# Focused Ultrasound Thermal Ablation and CD40 Agonism Reprograms Breast Tumor Immunity to Drive Regression and Memory

**DOI:** 10.64898/2026.03.02.708396

**Authors:** Zehra E. F. Demir, AeRyon Kim, Beyzanur G. Ak, Micaiah S.J. Lee, Thomas Sherlock, Stefanyda O. Maslova, Andrew T. Thede, Matthew R. DeWitt, Melanie R. Rutkowski, Natasha D. Sheybani

**Affiliations:** Department of Biomedical Engineering, University of Virginia, Charlottesville, VA; Department of Pathology, University of Virginia, Charlottesville, VA; Focused Ultrasound Immuno-Oncology Center, University of Virginia, Charlottesville, VA; Department of Microbiology, Immunology, and Cancer Biology, University of Virginia, Charlottesville, VA; Carter Immunology Center, University of Virginia, Charlottesville, VA; Department of Radiology & Medical Imaging, University of Virginia, Charlottesville, VA

## Abstract

Focused ultrasound thermal ablation (T-FUS) is a clinically accessible, non-invasive modality capable of inducing rapid tumor cytoreduction while mobilizing early immunologic danger signals. However, its capacity to synergize with potent co-stimulatory immunotherapies in breast cancer (BC) remains undefined. Here, we demonstrate that subtotal T-FUS cooperates with CD40 agonism to elicit durable, T cell-dependent tumor control across four immunologically and hormonally distinct murine BC models. Partial thermal ablation triggered canonical immunogenic cell-death signatures and acute remodeling of intratumoral myeloid populations, while expanding circulating CD4^+^ and CD8^+^ T cells. When layered onto this immunogenic milieu, αCD40 markedly constrained tumor outgrowth, yielding significant reductions in tumor burden across all models and complete tumor eradication in 33% of E0771 tumors, with additional complete responses in BRPKP110 and EMT6. Efficacy required both CD4^+^ and CD8^+^ T cells, and complete responders mounted robust systemic immunity, rejecting contralateral tumor rechallenge with 100% protection and displaying persistent effector-memory T cell activation. Together, these findings establish T-FUS as an immune-potentiating partner for CD40 agonism, capable of driving durable, robust BC regression and immunological memory. This work positions T-FUS+CD40 agonism as a clinically scalable, in situ vaccination-like strategy with potential to benefit breast cancers, including luminal subtypes, that remain largely refractory to immune checkpoint blockade.

## INTRODUCTION

Breast cancer (BC) has long been among the most commonly diagnosed cancers worldwide and remains a leading cause of cancer-related deaths in women^1–4^. Across both sexes, it ranks among the top cancers by incidence and among the leading causes of cancer mortality^5,6^. Recently, BC screening guidance has shifted, with the U.S. Preventive Services Task Force now recommending biennial mammography a decade earlier, starting at age 40^7^. Against this backdrop of rising BC incidence in younger women, priorities are increasingly focused on treatment approaches that maintain efficacy while reducing cumulative toxicity and long-term morbidity. However, current standards of care for BC still impose substantial patient burden. Surgery is inherently invasive and can entail prolonged convalescence (particularly in older patients) as well as breast deformity and suboptimal cosmesis. Systemic therapies such as chemotherapy can be dose-limited by hematologic toxicity, and both chemotherapy and radiation can carry short- and long-term adverse effects, including potential impacts on fertility - an especially salient consideration for younger patients^8,9^.

Less invasive, organ-sparing focal interventions have the potential to reshape BC treatment by reducing procedural morbidity and enabling unique opportunities for therapeutic synergy. Focused ultrasound (FUS) provides an incisionless modality for spatially precise thermal ablation (T-FUS)^10–12^ with a clinical track record in the treatment of benign^13,14^ and malignant^15,16^ breast tumors, and is now being explored as an adjunct to immunotherapies (ITx) (NCT04796220, NCT03237572)^15–17^. Preclinical and early clinical studies suggest that T-FUS can induce immunogenic tumor injury, including increased damage-associated molecular patterns (DAMPs; alarmins), upregulation of heat shock proteins, release of tumor-associated antigens, recruitment of antigen-presenting cells (APCs), and heightened local and systemic CD8+ and CD4+ T cell responses against solid tumors^12,15,18,19^. Accordingly, a key next step is to define how T-FUS can be leveraged to broaden and deepen ITx responses - particularly in BC, where this potential has yet to be systematically evaluated across distinct subtypes and immunophenotypes.

Clinical interest in ITx for BC continues to expand^20^, with PD-1/PD-L1 blockade improving outcomes in subsets of triple-negative disease^21^. However, benefit remains subtype-restricted and inconsistent across BC more broadly^22,23^, motivating combination strategies designed to expand and improve responses^20,24^. Given the marked heterogeneity of immune contexture across BC subtypes - including differences in baseline tumor-infiltrating lymphocytes^25^ - there is increasing interest in approaches that enhance priming rather than relying solely on reinvigoration of pre-existing T cell responses^24^. Among the most clinically advanced co-stimulatory strategies is CD40 agonism. CD40 (a TNF receptor superfamily member) is expressed predominantly on APCs, including dendritic cells, B cells, and macrophages. CD40 engagement promotes APC activation and licensing to support effective T cell priming^26–28^. Yet, despite encouraging activity in select settings, CD40 agonists have shown limited efficacy as monotherapies in solid tumors^26,29^, underscoring the need for rational partner interventions that increase immunogenic antigen availability and productive antigen presentation. In this context, T-FUS represents a mechanistically aligned modality to pair with CD40 agonism^24,30^.

Building on timely recent clinical evidence that an Fc-engineered CD40 agonist antibody (2141-V11) can induce tumor regression, including a complete response in BC^31^, we test the hypothesis that CD40 agonism cooperates with T-FUS to drive durable anti-tumor immunity in BC. Using a panel of murine models spanning diverse molecular subtypes and immunophenotypes, we show that combining αCD40 with a clinically aligned partial T-FUS regimen (NCT03237572) yields robust tumor control and survival benefit across multiple models. Mechanistically, therapeutic activity is T cell-dependent and can produce complete responses accompanied by evidence of immune memory. Collectively, these findings position T-FUS plus CD40 agonism as a clinically accessible combination strategy with the potential to extend ITx benefit across BC subtypes, including luminal tumors that are typically less responsive to classic checkpoint blockade^22,32^.

## RESULTS

### Subtotal FUS thermal ablation results in clear zones of apoptotic and necrotic tissue destruction, concomitant with local upregulation of canonical thermal stress signatures

First, we developed experimental methods and procedures to achieve partial thermal ablation of BC tumors, consistent with partial T-FUS benchmarks established in ongoing FUS breast immuno-oncology clinical trials (NCT04796220, NCT03237572). Briefly, we used a custom-ultrasound guided FUS system equipped with four therapeutic transducers driven at 3.78 MHz in continuous wave mode to elicit thermal energy deposition; focal temperatures were three dimensionally modeled *in silico* (Fig. 1A, Supplementary Fig. 1A-C). Tumors were acoustically coupled via degassed water bath and positioned directly over an axially co-registered ultrasound imaging probe, enabling real-time B-mode ultrasound guidance for tumor localization and targeting (Fig. 1B). Widely used in clinical applications, the appearance of hyperechoic marks in sonography are often used as an indicator of ablation^33–36^. During or immediately following T-FUS treatment, hyperechoic signatures were evident on B-mode (Fig. 1C). Strikingly, this varied from model to model, presumably on the basis of tissue stiffness, vascularity, or other intertumoral heterogeneities^37–39^. Among the E0771, BRPKP110, EMT6, and 4T1 models, 78.6%, 88.9%, 31.3%, and 94.4% of the mice were reported as a grade 3, respectively (Supplementary Table 1). Amongst these distinct BC models, we observed clear hyperechoic signals within all mice, with a majority (91.5%) at grade 2 or 3 (Fig. 1D, Supplementary Table 1).

**Figure 1:**
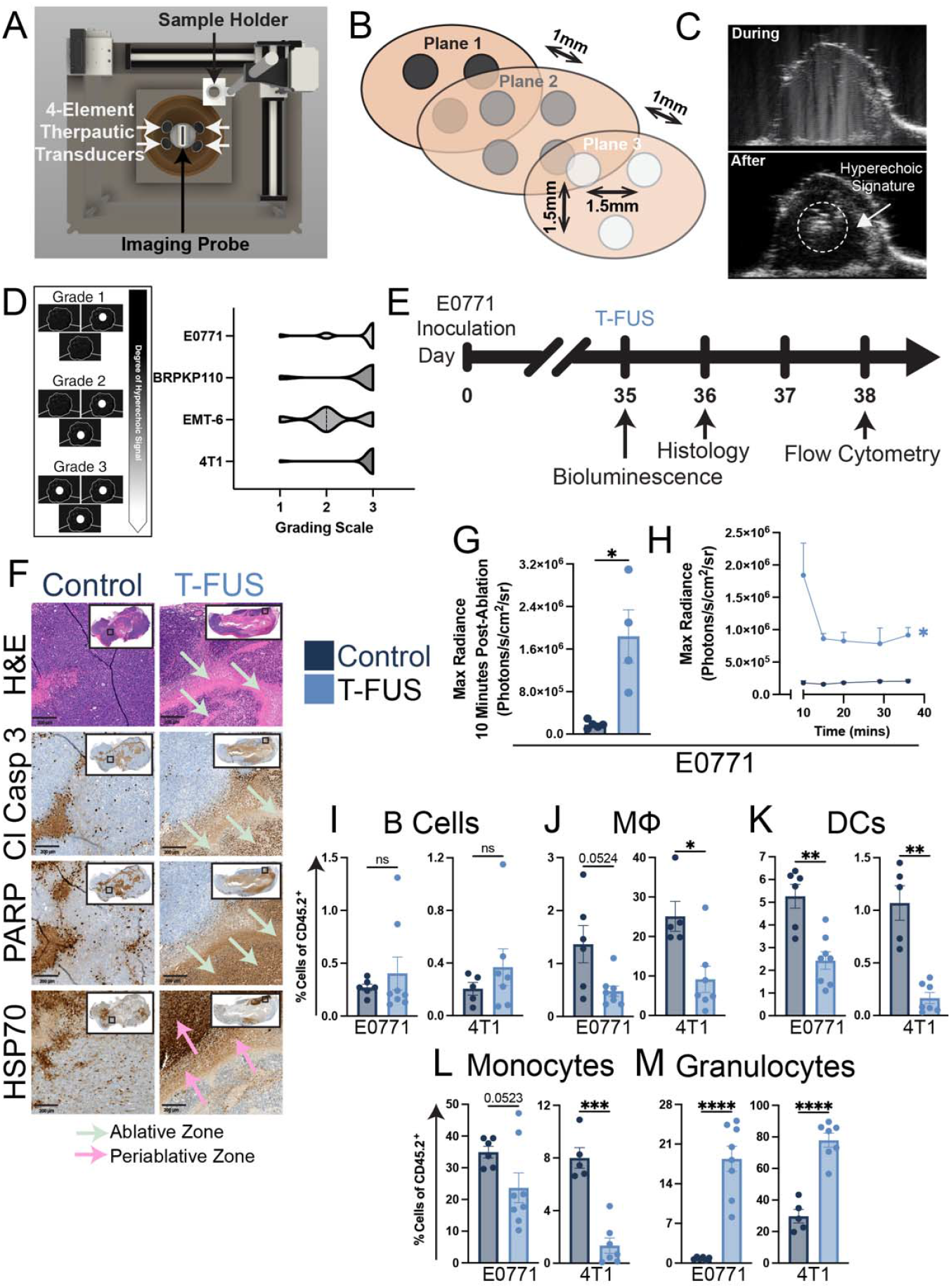
Partial thermal ablation induces stress signatures and acutely reshapes the tumor immune landscape. (A) CAD rendering of custom ultrasound-guided FUS system consisting of a 4-element 3.78 MHz transducer axially co-registered with 15 MHz MicroScan linear ultrasound imaging array. (B) Representative partial ablation scheme and (C) B-mode ultrasound images of E0771 tumor either before/during (top) or after (bottom) T-FUS treatment, with hyperechoic signature present after treatment (arrow). (D) Custom grading scale of presence of hyperechoic signal post-T-FUS treatment across all murine breast cancer models. (E) Overview of experimental timeline. (F) Representative H&E staining, along with cleaved caspase 3, PARP, and HSP70 staining, was performed on control and T-FUS-treated E0771 tumors 24 hours post-treatment. (G) *In vivo* surrogate measurements of ATP levels via bioluminescence at 10 minutes after T-FUS. (H) Temporal radiance curve (proxying relative ATP levels) for a sham vs. tumor volume-matched T-FUS mice (E0771 model). Significance assessed by Wilcoxon rank-sum test. **p=0.0028. H-L) (I) Percentage of B cells (CD19^+^, MHCII^+^), (J) macrophages (MФs; F480^+^, Ly6C^-^), (K) cDCs (CD11c^+^, MHCII^+^), (L) monocytes (F480^+^, Ly6C^+^), and (M) granulocytes (CD11b^+^, Ly6G^+^) out of CD45+ cells in the tumor. Significance assessed by Welch’s T-test.*p < 0.05, **p < 0.01, and ***p < 0.001 vs. control, respectively.

E0771 tumors were evaluated histologically one day post-treatment (Fig. 1E). Hematoxylin and eosin (H&E) staining of T-FUS treated tumors showed clear evidence of coagulative necrosis in the ablative zone, with patterning being distinct from tumor progression-associated necrosis in controls. In contrast with controls, ablative zones displayed higher levels and distinct patterning of apoptosis, as reflected by cleaved caspase 3 (Cl Casp 3) and poly(ADP-ribose) polymerase (PARP) staining (Fig. 1F). Furthermore, upregulation of heat shock protein (HSP70) staining was evident in the so-called “periablative zone”, i.e. regions neighboring those bearing signatures of coagulative necrosis and apoptosis – consistent with the notion of a transition zone from heat-mediated cell death to damage from ablative and periablative regions, respectively (Fig. 1F).

Release of adenosine triphosphate (ATP) is considered a hallmark of immunogenic cell death (ICD), acting as a key DAMP^40^. T-FUS acutely conferred an ∼146-fold increase in average luminescence relative to sham conditions in 4T1 cells *in vitro* (Supplementary Fig. 2A). Translated to orthotopic E0771 BCs *in vivo*, we observed that T-FUS conferred a ∼10.3-fold increase in average maximum radiance, a proxy for ATP release, at 10 minutes post-treatment (Fig. 1G). Longitudinal bioluminescence imaging revealed superlative intra/peritumoral ATP levels in T-FUS-treated tumors compared to the volume-matched shams, and interestingly, this trend was sustained over the 40-minute imaging period (Fig. 1H).

Three days following T-FUS, tumors were excised for immunological characterization by multispectral flow cytometry. T-FUS, analogously deployed in E0771 and 4T1 tumors, did not confer changes to intratumoral B cells (CD19^+^, MHCII^+^; Fig. 1I). However, the percentage of macrophages (MФs; F4/80^+^, Ly6C^+^), conventional dendritic cells (cDCs; CD11c^+^, MHCII^+^), and monocytes (Ly6C^+^, F4/80^-^) was significantly decreased following ablation (Fig. 1J-L). Conversely, observed across both BC models, there was an acute enrichment in local granulocytes (Ly6G^+^, CD11b^+^), likely neutrophils and/or granulocytic myeloid-derived suppressor cells (G-MDSCs) (Fig. 1M). Thus, subtotal T-FUS produces a spatially patterned thermal injury - coagulative necrosis and apoptosis with a periablative heat-stress border - coupled to rapid ATP release and an acute remodeling of the intratumoral myeloid compartment toward granulocytic predominance.

### Subtotal FUS thermal ablation promotes a systemic T cell response

We also investigated shifts in the T lymphocyte population in E0771 mice 3 days post-treatment (Fig. 2A). As anticipated, there was a significant decrease in the absolute number per gram and percentage of CD8^+^ and CD4^+^ T cell (including Treg) subsets in the tumor (Fig. 2B-G). While we observed an overall decrease in intratumoral T cell numbers, T-FUS-treated tumors exhibited a significantly higher CD8:CD4 T cell ratio relative to control indicating a compositional shift toward CD8-skewed intratumoral T cell balance 3 days post-treatment (Fig. 2H). At this time point, we did not anticipate T-FUS to augment intratumoral T cells, but existing literature suggesting that T-FUS can induce a systemic T cell response^17,41^ prompted us to postulate that T-FUS may elaborate systemic T cell signatures. Indeed, in the tumor draining lymph node (TDLN), we observed strong trends toward enriched CD8^+^ and CD4^+^ T cell numbers, while the percentage of these cells among total lymphocytes remained unchanged (Fig. 2I-L). In tandem, we sampled the circulating immune cell repertoire via terminal cardiac bleed. T-FUS treatment significantly increased the number of CD8^+^ T cells, while driving a strong trend toward increased CD4^+^ T cells (∼2-fold) (Fig. 2M-O). Moreover, T-FUS drove a significant increase in percentage of both CD8^+^ and CD4^+^ T cells (Fig. 2P-Q). Together, these results support that beyond its function as a local debulking modality, T-FUS functions as a systemic immunologic stimulus, yielding favorable shifts in local T cell ratio and elevating T cell representation in TDLN and blood.

**Figure 2:**
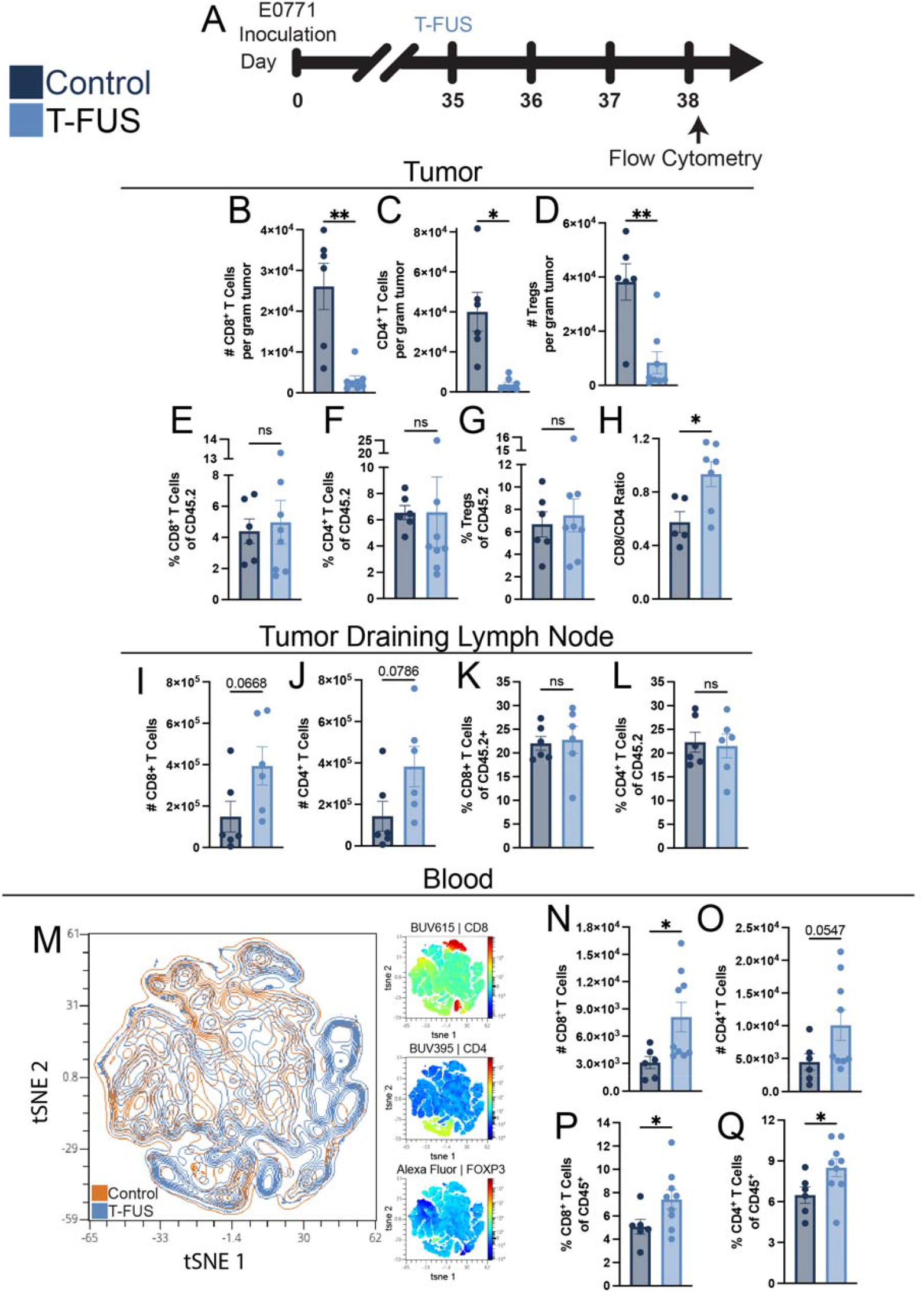
Partial thermal ablation acutely remodels T lymphocyte representation across tumor and periphery. (A) Experimental timeline for T-FUS treatment and downstream analysis. (B-G) Absolute number per gram of tumor and percentage of intratumoral CD8^+^, CD4^+^, and Foxp3^+^ (regulatory) T cells. (H) Ratio of intratumoral CD8^+^ and CD4^+^ T cells. (I-L) Number and percentage of T lymphocytes in the tumor draining lymph node (TDLN). (M) Multigraph color mapping of tSNE plot on circulating CD45^+^ cells. (N-O) Absolute number and (P-Q) percentage of circulating CD8^+^ and CD4^+^ T cells in whole blood. Significance assessed by Welch’s T-test.*p < 0.05, **p < 0.01 vs. control.

### Subtotal FUS thermal ablation in combination with a CD40 agonist constraints tumor outgrowth and promotes survival - conferring complete responses in multiple breast cancer models

Consistent with preclinical^11,18,19^ and clinical^15,16,42,43^ findings, our results support the role of partial thermal ablation as an immunomodulator. Accordingly, we sought to evaluate whether coupling of CD40 agonism would act in cooperation, enabling a promising combinatorial strategy for BC. To evaluate the efficacy of T-FUS and αCD40, across all BC models, we initiated treatment with i.p. administrations of αCD40 (100 *μ*g/mouse) every 3 days once tumors reached approximately 60mm^3^ (Fig. 3A). We utilized multiple different murine models reflecting the diversity of human BC subtypes in their hormone receptor status, immune mosaic, and metastatic proclivity. These include basal-like triple negative BC models including 4T1 and EMT6 (Balb/c background), as well as hormone receptor positive BC models including BRPKp110 and E0771 (C57BL/6 background) – which recapitulate Luminal A-like^44^ and Luminal B-like^45^ subtypes, respectively.

**Figure 3:**
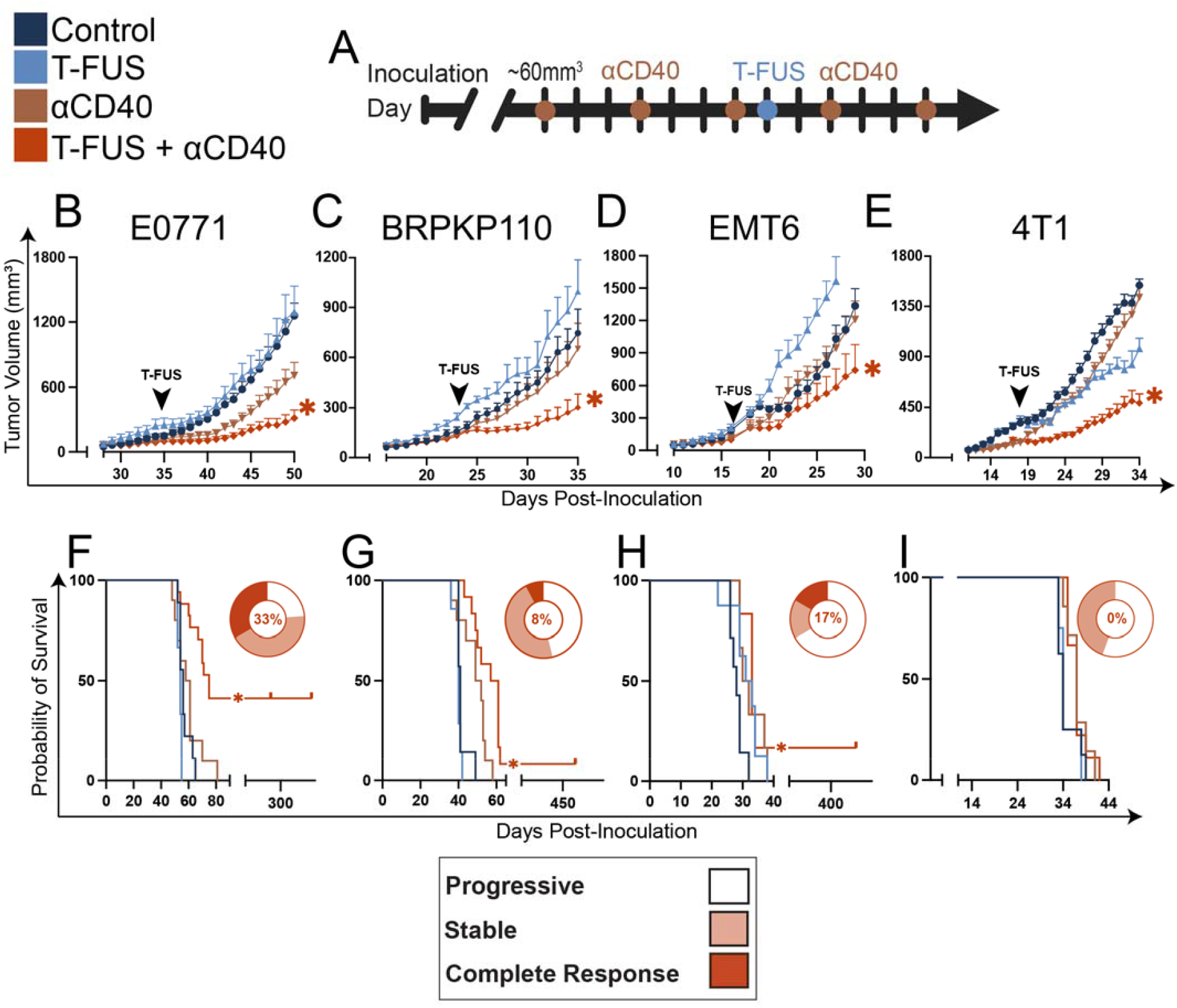
Thermal ablation and CD40 agonism cooperate to constrain distinct breast cancer subtypes and extend survival. (A) General experimental timeline for T-FUS+αCD40 paradigms across all murine breast cancer models. (B-E) Primary tumor outgrowth for E0771, BRPKP110, EMT-6, and 4T1 tumor-bearing mice, receiving IgG, T-FUS monotherapy, αCD40 monotherapy, or T-FUS+ αCD40. Significance assessed by 2-way ANOVA, followed by Holm-Sidak multiple comparison correction. *E0771:* p<0.05 vs all other groups (specifically, IgG (n=15) vs T-FUS+αCD40 (n=21): p = <0.0001; T-FUS (n=7) vs T-FUS+αCD40: p =0.0114; αCD40 (n=15) vs T-FUS+αCD40: p = 0.0116). *BRPKP110*: p<0.05 vs all other groups (specifically, IgG (n=4) vs T-FUS+αCD40 (n=3): p = 0.0134; αCD40 (n=5) vs T-FUS+αCD40: p = 0.0354). *EMT-6*: p<0.05 vs all other groups (specifically, IgG (n=7) vs T-FUS+αCD40 (n=6): p = 0.0047; T-FUS (n=8) vs T-FUS+αCD40: p = 0.0862; αCD40 (n=6) vs T-FUS+αCD40: p = 0.0437). *4T1*: p<0.05 vs all other groups (specifically, IgG (n=8) vs T-FUS+αCD40 (n=9): p<0.0001; T-FUS (n=4) vs T-FUS+αCD40: p = 0.0045; αCD40 (n=7) vs T-FUS+αCD40: p <0.0001). (F-I) Kaplan-Meier curve depicting overall survival of E0771, BRPKP110, EMT6, and 4T1 tumor models in the combination paradigm, accompanied by a pie chart illustrating the distribution of mice with progressive disease, stable disease, and complete responses. Percentages represent the proportion of complete responders within each respective group. Significance assessed by log-rank (Mantel-Cox) test. *E0771*: p<0.05 vs all other groups (specifically, IgG vs T-FUS+αCD40: p <0.0001; T-FUS vs T-FUS+αCD40: p = 0.0005; αCD40 vs T-FUS+αCD40: p = 0.0019). *BRPKP110*: p<0.05 vs all other groups (specifically, IgG vs T-FUS+αCD40: p = 0.0126; αCD40 vs T-FUS+αCD40: p = 0.0439). *EMT-6*: p<0.05 vs all other groups (specifically, IgG vs T-FUS+αCD40: p = 0.0019; T-FUS vs T-FUS+αCD40: p = 0.417; αCD40 vs T-FUS+αCD40: p >0.9999). *4T1:* p<0.05 vs all other groups (specifically, IgG vs T-FUS+αCD40: p = 0.0758; T-FUS vs T-FUS+αCD40: p = 0.0732; αCD40 vs T-FUS+αCD40: p = 0.8525).

Across all BC models (E0771, BRPKP110, EMT6, 4T1), neither monotherapy offered significant benefit whereas the combination treatment group offered the most superlative constraint of tumor outgrowth relative to controls (Fig. 3B-E). By a week following the conclusion of treatments, E0771, BRPKP110, EMT6, and 4T1 tumors exposed to T-FUS+αCD40 combination saw nearly 3.5X, 2.5X, 2.3X, and 3.1X reduction in the average tumor volume compared to IgG control, respectively. Area under the curve (AUC) analysis revealed significant reduction of cumulative tumor burden in E0771 and 4T1, with a striking 1.7-fold and 2-fold reduction in AUC compared to αCD40 monotherapy, respectively (Supplementary Fig. 2B-C). Serial 3D B-mode ultrasound volumetry of 4T1 tumors independently confirmed a significant reduction in tumor burden in the combination group versus all other groups, corroborating digital caliper measurements (Supplementary Fig. 2D-E).

Combination of T-FUS and CD40 agonism also promoted significant survival benefit across 3 of 4 models (E0771, BRPKP110, and EMT6), underscoring the robustness of the combination across BC subtypes but also the importance of subtype-specific considerations (Fig. 3F-I). E0771 tumor-bearing mice receiving T-FUS+αCD40 additionally saw the most impressive extension in overall survival, exhibiting ∼34% ∼39%, and 26% increases in median survival time relative to IgG, T-FUS, and *α*CD40 groups, respectively (Fig. 3F). Consistent with clinical response assessment frameworks, we assigned each mouse to a response category - progressive disease, stable disease, or complete response - to further probe response to T-FUS+αCD40 (Fig. 3F-I; wheel insets). Remarkably, E0771 mice treated with the combination displayed the highest proportion of stable and/or complete responses. In the combination treatment group exclusively, 33% of the mice demonstrated complete tumor regression (complete responders; CRs) (Fig. 3F). Furthermore, across two additional BC models – BRPKP110 and EMT6 – CRs were observed only in the T-FUS+αCD40 group, albeit to a lesser degree (Fig 3G-H). While 4T1 tumor-bearing mice did not receive explicit survival benefit, likely owing to onset of spontaneous pulmonary metastasis known to this model, ∼44% of mice experienced stabilization of primary disease (Fig. 3I). In aggregate, T-FUS+αCD40 consistently outperformed controls across all models tested, delivering the most robust suppression of tumor outgrowth, extending survival across multiple models, and inducing stable/complete responses with BC subtype-specific magnitude. In light of the particularly robust response observed in E0771, subsequent experiments focused on this model to enable deeper mechanistic dissection.

### αCD40 priming orchestrates potent local and system immune activation, enriching the abundance and maturity of APCs and T cells

Given the pronounced efficacy of the T-FUS+αCD40 regimen, we next sought to define how αCD40 priming reshapes the local and systemic immune landscape prior to ablation, thereby establishing the immune context in which T-FUS is delivered. We utilized multispectral flow cytometry to identify local and systemic changes in APCs in E0771 tumor-bearing mice (Fig. 4A). tSNE visualization revealed qualitative shifts in the APC population in TDLN, guiding subsequent analysis by conventional gating (Fig. 4B). Consistent with findings in the literature^26–28^, αCD40 priming increased the absolute number and percentage of APCs including B cells, MФs, and cDCs (Fig. 4C-H). Additionally, the geometric mean fluorescent intensity (GMFI) of CD86, a canonical co-stimulatory marker of APC activation, on these cells was also significantly elevated with CD40 agonism (Fig. 4I-K). Moreover, the percentage of CD86 on these cells was also increased in the TDLN (Fig. 4L-N). Upon further disaggregation of the cDC population in the TDLN, we observed an increase in both the number and percent of cDC1s (CD11b^+^, XCR1^+^) and cDC2s (CD11b^+^, XCR1^-^, SIRPα^+^) (Supplementary Fig. 3A-D). Additionally, we observed an increase in the percentage of CD86^+^ cDC1s and GMFI of CD86 on cDC1s and cDC2s in the TLDN (Supplementary Fig. 3E-I). Beyond the TDLN, the global impacts of αCD40 priming were evident through shifts in the splenic and contralateral lymph node (cLN) B cell and MФ populations (Supplementary Fig. 4A, H), which displayed significantly elevated absolute numbers and maturity (Supplementary Fig. 4B-G, I-N).

**Figure 4:**
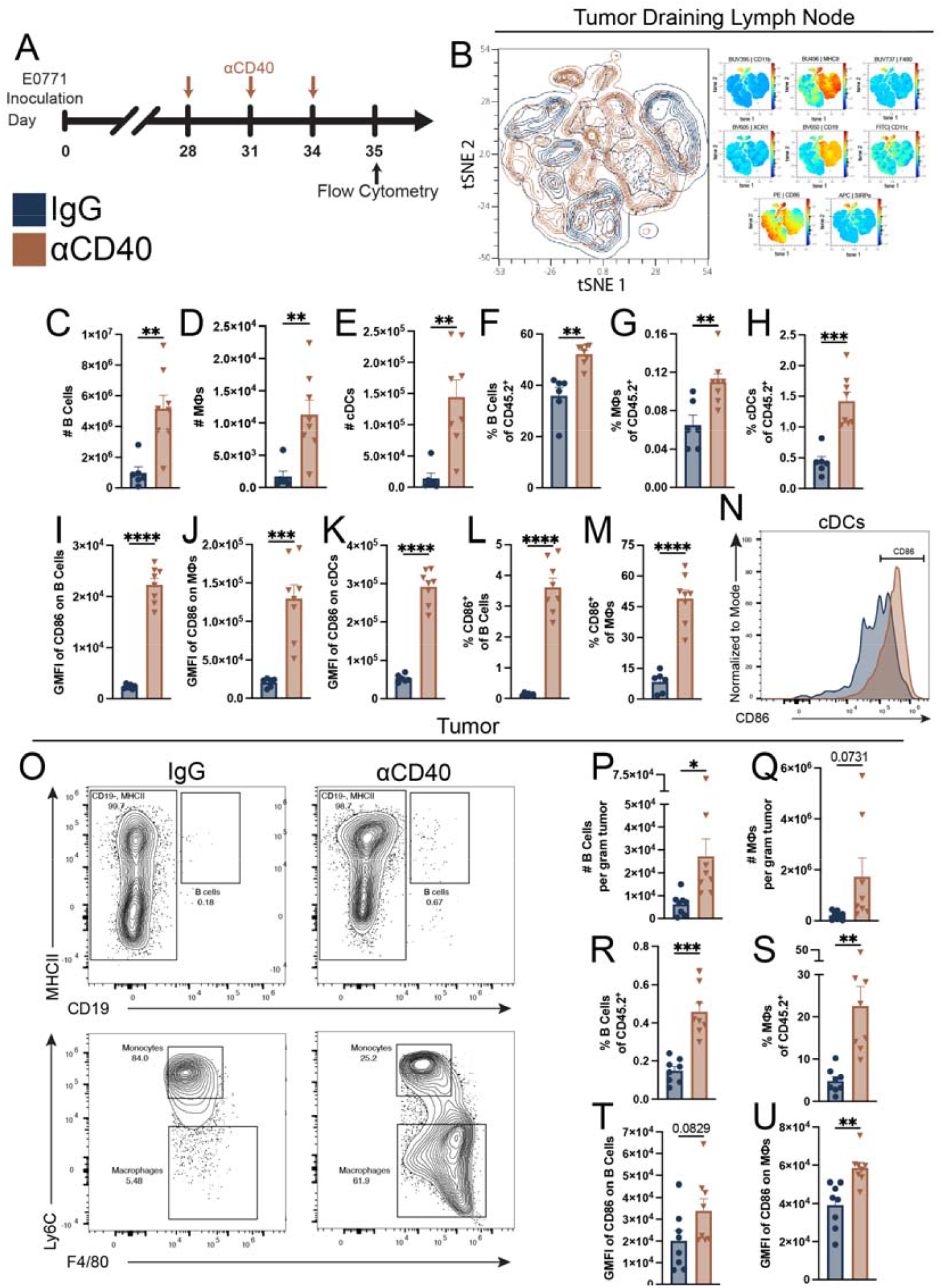
aCD40 priming prior to T-FUS activates antigen-presenting cells locally and systemically. (A) Experimental timeline for CD40 priming. (B) Multigraph color mapping of tSNE plot on CD45^+^ immune cells in the TDLN. (C-H) Absolute number and percentage of B cells, MФs, and cDCs in the TDLN. (I-K) The GMFI of CD86 and (L-N) percentage of CD86^+^ B cells, MФs, and cDCs in the TDLN. (N) Histogram representing the percentage of CD86^+^ cDCs in the TDLN. (O) Representative dot plot of intratumoral B cell, monocytes, and MФs. (P-U) Absolute number per gram and percentage of intratumoral B cells and MФs, as well as GMFI of CD86 on these subsets. Significance assessed by Welch’s T-test.*p < 0.05, **p < 0.01, and ***p < 0.001 vs. IgG control.

We co-examined the local effects of tumor priming with CD40 agonism. While neither the absolute number per gram of tumor nor the percentage of cDCs changed, we observed an increase in the number and percentage of B cell and MФs (Fig. 4O-S, Supplementary Fig. 3J-K). Additionally, we observed a trend in greater CD86 GMFI on B cells, which was significant on MФs (Fig. 4T-U). In addition to augmenting intratumoral APCs, CD40 priming elicited shifts in the intratumoral T lymphocyte landscape (Supplementary Fig. 5A-B), specifically yielding significantly increased absolute number and percentage of intratumoral CD8^+^ T cells (Supplementary Fig. 5C, F); increased percentage of CD4^+^ T cells (Supplementary Fig. 5D,G); and significantly decreased percentage of Tregs (Supplementary Fig. 5E,H). Following αCD40 priming, both CD8^+^ and CD4^+^ T cells upregulated CD40L, exhibiting >2-fold higher CD40L GMFI, consistent with an activated phenotype and enhanced capacity for CD40-dependent costimulatory signaling (Supplementary Fig. 5I-J). To begin distinguishing whether αCD40-associated shifts in T-cell abundance reflected local proliferation versus redistribution/trafficking, we also assessed Ki-67 on CD8^+^ and CD4^+^ T cells. T cells exhibited a significant increase in the percentage of Ki-67^+^ cells and in Ki-67 GMFI with CD40 agonism (Supplementary Fig. 5K-O). Taken together, these data indicate that αCD40 priming drives broad APC activation across compartments and markedly increases intratumoral T-cell abundance and proliferative state just prior to T-FUS.

### Complete and non-complete responders to TFUS+αCD40 are distinguished by their T cell signatures

A subset of mice (CRs) achieved complete tumor eradication in the combination arm (Fig. 5B-D). In an effort to identify immunologic correlates of durable tumor control, we leveraged the coexistence of these complete and “non-complete” responders (non-CRs) and profiled the T-cell landscape at a chronic endpoint (Fig. 5A). We dissected the immunological repertoire in tumors (when available) and secondary lymphoid organs via multispectral flow cytometry. Of note, no tumor data could be collected from CRs since their tumors had been entirely eliminated by the time of assessment. The tumors of combination-recipient non-CRs were compared to monotherapy and control counterparts. Despite appreciable tumor growth control in non-CRs, we observed that intratumoral T cell representation was indistinguishable from mice receiving αCD40 alone (Fig. 5E-H). Both groups equivalently exhibited a trend towards increased CD8^+^ T cell numbers and significant elevation in CD8^+^ T cell percentage compared to T-FUS and IgG controls (Fig. 5E-G). Intratumoral CD4^+^ T cell representation was largely unchanged at this time point (Fig. 5F,H). These data suggest that, in non-CRs, T-FUS may not have sufficiently augmented intratumoral T cell signatures beyond those elicited by αCD40 alone, potentially contributing to incomplete tumor control.

**Figure 5:**
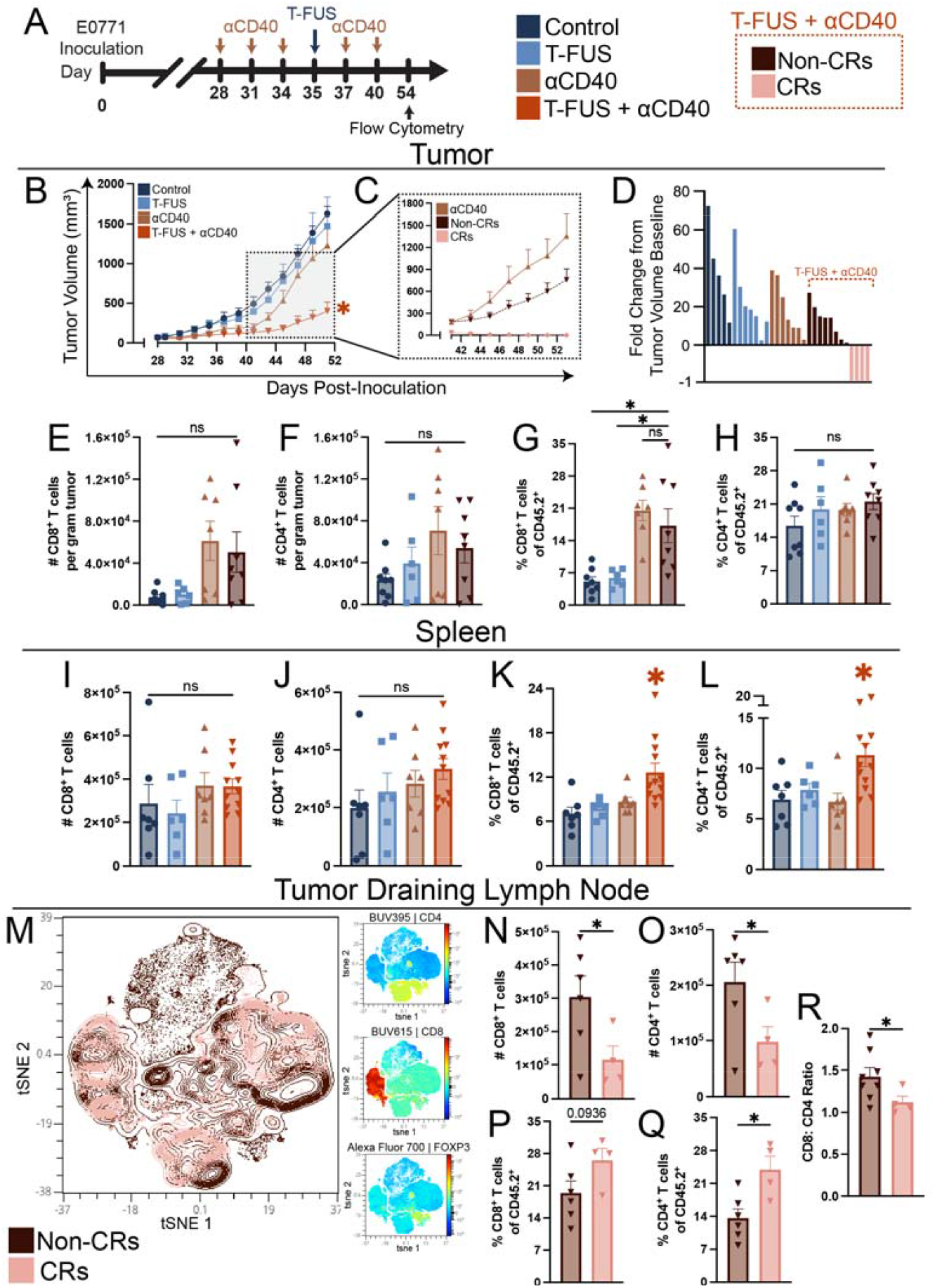
Combination of T-FUS and aCD40 yields complete responders, which bear T lymphocyte signatures distinct from non-complete responders. (A) Experimental timeline of T-FUS+αCD40 dosing and harvest for flow cytometry analysis. (B) Average tumor outgrowth of E0771 tumor-bearing mice. Significance assessed by 2-way ANOVA, followed by Holm-Sidak multiple comparison correction. *E0771:* p<0.05 vs all other groups (specifically, IgG (n=5) vs T-FUS+αCD40 (n=12): p = <0.0001; T-FUS (n=7) vs T-FUS+αCD40: p =0.0843; αCD40 (n=7) vs T-FUS+αCD40: p = 0.0517). (C) Focused view of tumor outgrowth at selected timepoints (day 42-53) highlighting differences between T-FUS+αCD40 non-CRs and CRs, versus αCD40 monotherapy. (D) Waterfall plot depicting CRs in the combination treatment group. (E-H) Absolute number per gram tumor and percentage of CD8^+^ and CD4^+^ T cells in the tumor and (I-L) spleen. M) tSNE plot on CD45^+^ cells in the TDLN. (N-Q) Absolute number and percentage of CD8^+^ and CD4^+^ T cells in the TDLN. (R) Ratio between CD8^+^ and CD4^+^ T cells in the TDLN. Significance assessed using a Welch ANOVA or Welch’s T test. *p < 0.05, **p < 0.01 vs. T-FUS+αCD40 CRs.

We concomitantly evaluated spleens and TDLNs. While splenic T cell numbers did not change appreciably (Fig. 5I-J), we observed a significant increase in their percentages across all combination recipients, irrespective of their response level (Fig. 5K-L). Contrastingly, T cell signatures within TDLNs were distinguishable by response level. Specifically, CRs exhibited reduced TDLN T cell cellularity but increased relative representation of T cells (particularly CD4^+^), resulting in a lower CD8:CD4 ratio - consistent with a contracted or resolved TDLN state after tumor eradication; non-CRs retained relatively expanded, CD8^+^-skewed TDLNs consistent with persistent disease (Fig. 5M-R). These findings indicate that T-FUS+αCD40 elicits measurable systemic T-cell shifts while revealing response-associated differences in the draining lymph node T cell landscape.

### Protective effect of T-FUS+αCD40 is T cell-dependent

Given the robust efficacy of T-FUS+αCD40 and the systemic immune activation observed with αCD40 priming, we hypothesized that therapeutic benefit from the combinatorial regimen requires T cells. To test this, mice receiving T-FUS+αCD40 were depleted of CD4^+^ T cells, CD8^+^ T cells, or both during the post-treatment effector window (D41-59; Fig. 6A), with efficient depletion confirmed in peripheral blood by flow cytometry (Fig. 6B). Strikingly, loss of either CD4^+^ or CD8^+^ T cells abrogated tumor control. Compared to isotype-treated controls, mean tumor volumes were ∼2.8-3-fold higher in CD4-depleted, CD8-depleted, and dual-depleted groups (Fig. 6C). Concordantly, T cell depletion significantly reduced overall survival (Fig. 6D), eliminating the durable protection observed in isotype-treated controls - wherein a subset of mice saw complete response. Together, these findings demonstrate that the protective effect of T-FUS+αCD40 is T cell-dependent, with both CD4^+^ and CD8^+^ compartments contributing to therapeutic efficacy.

**Figure 6:**
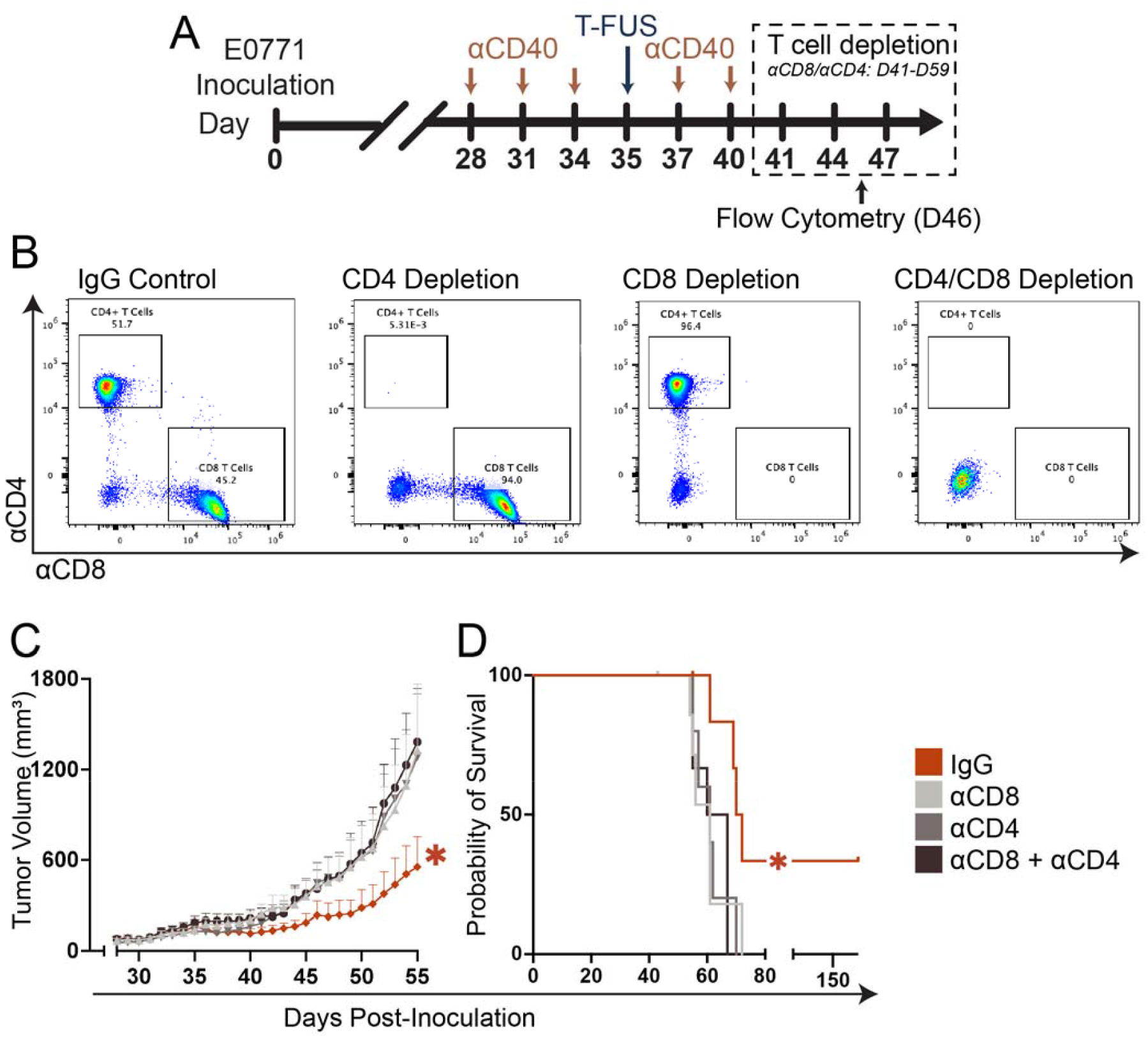
Protective effect of focused ultrasound thermal ablation and CD40 agonism is dependent on T cells. (A) Overview of timeline for T cell depletion conducted on T-FUS+αCD40 background. (B) Representative flow plots confirming effective depletion of CD8^+^/CD4^+^, CD8^+^, or CD4^+^ T cells. (C) Average tumor outgrowth on T-FUS+αCD40 (WT) background - with or without αCD4, αCD8, or αCD4 + αCD8 depletion. Significance assessed by 2-way ANOVA, followed by Holm-Sidak multiple comparison correction. *p<0.05 vs all other groups (specifically, αCD8 (n=6) vs IgG (n=5): p = 0.0005; αCD4 (n=5) vs IgG: p =0.0009; αCD4/αCD8 (n=6) vs IgG : p = 0.0003). (D) Kaplan-Meier curve depicting impact of T cell depletion on overall survival in T-FUS+αCD40-recipient E0771 mice. Significance assessed by log-rank (Mantel-Cox) test. p<0.05 vs all other groups (specifically, αCD8 (n=6) vs IgG (n=5): p = 0.0272; αCD4 (n=5) vs IgG: p = 0.0382; αCD4/αCD8 (n=6) vs IgG : p = 0.0098).

### Complete responders reject contralateral re-challenge and exhibit systemic, functional T-cell memory

CRs in the T-FUS+αCD40 group remained tumor-free for >150 days following treatment, and upon contralateral re-challenge with 1e6 E0771 cells (2x the primary inoculum; Fig. 7A), uniformly rejected tumor outgrowth compared to naïve, age-matched controls. This was reflected in complete protection from tumor establishment and 100% survival in the CR cohort, whereas controls developed tumor burden and succumbed to disease irrespective of age (Fig. 7B-D). To define immune correlates of this durable protection, we serially profiled peripheral blood by multispectral flow cytometry following re-challenge. CRs displayed a significant increase in the percentage of antigen-experienced/effector memory CD4^+^ T cells (CD44^+^, CD62L^−^) compared to treatment-naïve controls (Fig. 7E-F). Although there was no observable change in the number or percentage of peripheral CD8^+^ and CD4^+^ T cells following rechallenge (Fig. 7G-J), CD40L (Fig. 7K-N) and Ki-67 (Fig. 7O-R) expression were significantly increased on both circulating T cell subsets. These observations are consistent with an activated, proliferative recall state and enhanced CD40–CD40L costimulatory signaling capacity that could reinforce APC engagement during protective immunity in CR mice. Furthermore, the CD4 compartment in CRs exhibited an enhanced effector cytokine signature, with increased representation of IFN-γ^+^ and IFN-γ^+^TNF-α^+^ antigen-experienced CD4^+^ T cells (Fig. 7S-W). Collectively, these findings support that durable cures achieved with T-FUS+αCD40 are associated with a systemic recall-type T-cell response marked by expanded antigen-experienced CD4^+^ memory, enhanced proliferation, and heightened effector function following re-challenge.

**Figure 7:**
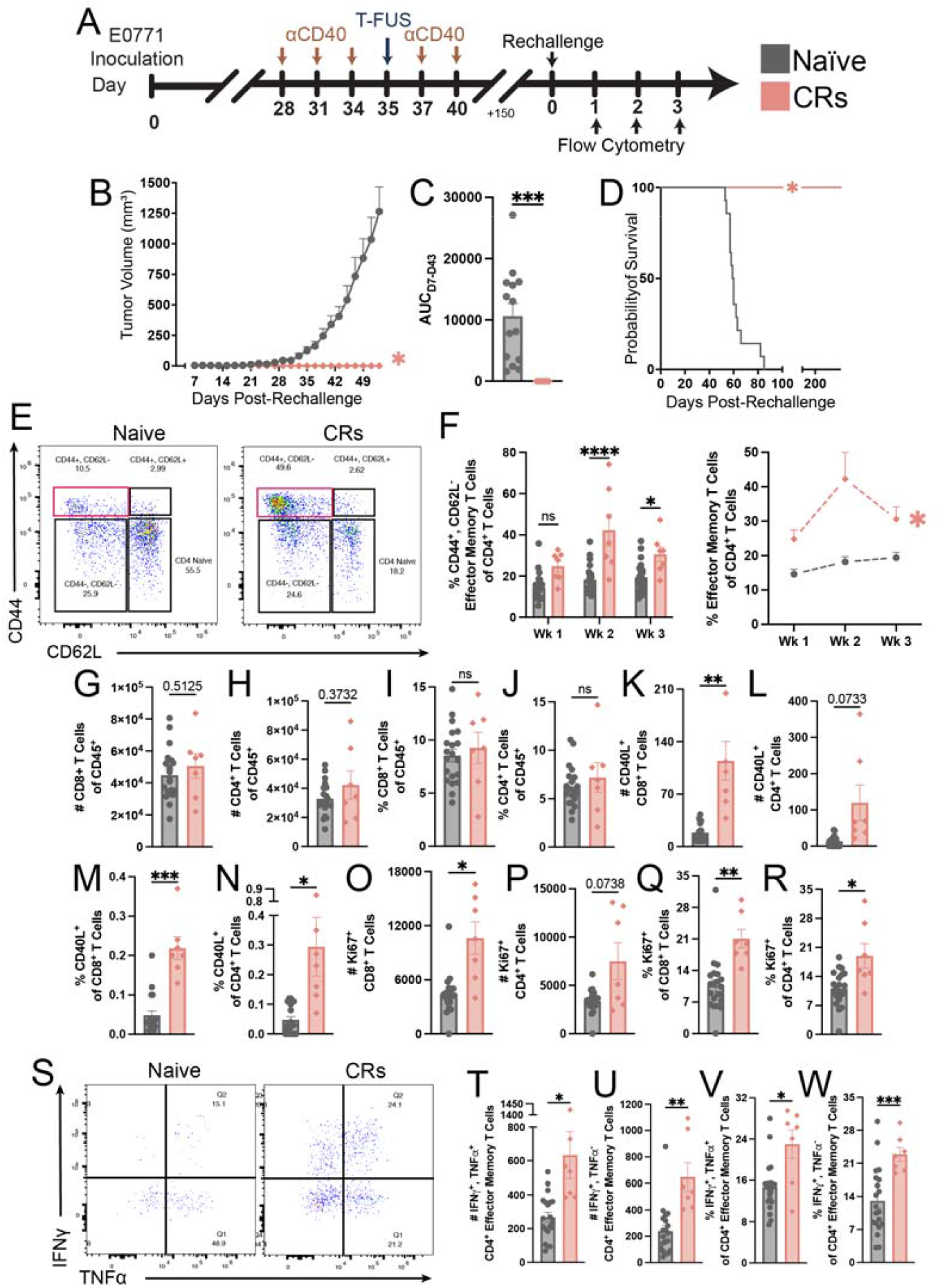
Complete responders to T-FUS + CD40 agonism reject tumor rechallenge and exhibit systemic immune memory. (A) Experimental timeline depicting the T-FUS+αCD40 dosing and rechallenge timeline. (B) Average E0771 tumor outgrowth in naïve, age-matched, WT mice and re-challenged CR mice. Significance assessed by 2-way ANOVA, followed by Holm-Sidak multiple comparison correction. *p<0.05 vs all other groups (specifically, naïve (n=14) vs CRs (n=7): p = 0.0007). (C) AUC analysis from day 7 to day 43. Significance assessed by Welch’s T test (specifically, naïve vs CRs: p = 0.0002). (D) Kaplan-Meier curve depicting 100% survival in mice rechallenged. (E) Representative flow plots of memory T cells in peripheral blood at week 2 post-rechallenge. (F) Percentage of effector memory CD4^+^ T cells (*CD*4+, *CD*62L− ) in peripheral blood at week 1-3 following rechallenge. (G-J) Absolute number and percentage of CD8^+^ and CD4^+^ T cells. (K-N) Percentage of CD40L^+^ as well as (O-R) Ki67^+^ CD8^+^ and CD4^+^ T cells week 1 post-rechallenge. (E) Representative dot plots of IFN-γ^+^, TNFα ^+^ CD4^+^ effector memory T cells following rechallenge in the naïve and CRs groups. (T-W) Number and percentage of *IFNγ*+, *TNFα*+ and *IFNγ*+, *TNFα*-CD4^+^ effector memory T cells in the blood 1 week post-rechallenge. Significance assessed using Welch’s T test. p < 0.05, **p < 0.01, and ***p < 0.001 vs. Naïve control.

## DISCUSSION

In this study, we establish subtotal focused ultrasound thermal ablation (T-FUS) as a potent partner for CD40 agonism in BC. To our knowledge, this is the first report to extend a T-FUS immuno-oncology paradigm across BC models that reflect the heterogeneity of molecular subtypes represented in human disease^46,47^. Across multiple genetically, hormonally, and immunologically distinct models^45,48,49^, T-FUS+αCD40 consistently outperformed either monotherapy, delivering the most robust constraint of tumor outgrowth and yielding complete responses in multiple settings. Immunotherapy responses in BC have largely been concentrated in triple negative settings^24^, whereas hormone receptor-positive luminal disease is comparatively immunologically “silent”^50,51^. Thus, a critical contribution of our study stems from the inclusion of E0771, a luminal B-like HR^+^/ ERα^-^ /ERβ^+45,48^, and BRPKp110, a luminal A-like HR^+^/ERα^+52^ syngeneic model, which effectively expands exploration into a disease context that dominates human BC^50^ but is to date understudied in FUS immuno-oncology. To our knowledge, BRPKp110 has not previously been evaluated in the setting of FUS. The breadth of efficacy that we observe with the combination of T-FUS and αCD40 - extending into luminal-like contexts - underscores the potential of subtotal T-FUS to dramatically expand the immunotherapy-responsive landscape of BC going forward. With these findings, we conclude that T-FUS+αCD40 is a cooperative strategy capable of translating local tumor injury into adaptive, systemic antitumor immunity and unveil insights for future tuning of this combination in a BC subtype-specific manner.

Our results support a working model in which αCD40 priming establishes a permissive antigen presentation landscape that can be productively leveraged by a single session of intervening subtotal ablation. Prior to T-FUS, αCD40 increased the abundance and activation state of APCs - particularly within the TDLN - consistent with enhanced priming capacity and APC licensing effects corroborated within other studies^48,53–55^. Moreover, a recent publication corroborates our findings in E0771 & BRPKp110, demonstrating that the administration of a CD40 agonist effectively induces the recruitment of cDC1s and cDCD2 within the TDLN, concurrently enriching trafficking of intratumoral T cells^48^. Indeed, this underscores the rationale behind alliance with T-FUS to induce a durable “vaccine-like” effect. T-FUS then triggered hallmark spatially patterned thermal denaturation, canonical thermal stress signatures (i.e. heat shock protein), and rapid alarmin (i.e. ATP) release that collectively drove appreciable acute shifts in the tumor-immune landscape^11,18,19,43^. Interestingly, across two BC models with fundamentally distinct vascular, stromal and immunological compositions^11,56^, T-FUS elicited convergent early immune remodeling – marked by significant reductions in intratumoral APCs (except B cells) and monocytes, as well as a pronounced granulocytic influx consistent with acute thermal injury^57–59^. Notably, this influx neither accelerated tumor progression nor was it deleterious to combinatorial benefit. While absolute intratumoral T-cell numbers contracted, their relative proportion was preserved acutely following T-FUS, coincident with early enrichment of CD4^+^ and CD8^+^ T-cell signatures in systemic compartments (TDLN and blood) – suggesting that T-FUS-driven cytoreduction is not inherently counterproductive to T-cell responses. Nevertheless, T-FUS monotherapy did not yield durable tumor control, suggesting that additional immune activation is required to yield sustained antitumor benefit.

It was only when T-FUS was combined with αCD40 that we observed significant primary growth control of BC tumors (4 of 4 models) and survival benefit (3 of 4 models). We observed the most durable control – and the highest proportion of CRs – in E0771 and more modest effects in BRPKP110, EMT6, and 4T1; this variability likely reflects differences in baseline immune contexture, metastatic propensity, host strain, and/or delivered thermal dose. Future studies should explore thermal dose, ablation volume, sonication pattern and density, and therapeutic sequencing to further define determinants of response.

We focused subsequent mechanistic studies on E0771 because it reproducibly yielded distinct response phenotypes (CRs vs non-CRs), enabling higher-resolution dissection of immune correlates associated with durable tumor control. We determined that therapeutic benefit and CRs conferred by the combination required adaptive immunity. Depletion of CD4^+^ and/or CD8^+^ T cells not only abrogated tumor control and survival benefit but also abolished complete responses - supporting a central role for T cell effector function in the combination paradigm. CRs rejected tumor re-challenge delivered at twice the original inoculum and exhibited distinct circulating T cell signatures that were consistent with functional systemic antitumor immune memory. Specifically, they exhibited a clear recall phenotype after re-challenge, defined by CD4^+^ T cells enriched for markers of antigen experience, proliferation, CD40L-mediated costimulatory potential, and effector function. While this is strongly consistent with a tumor-directed, memory T-cell response underscoring rechallenge rejection, tumor antigen specificity was not explicitly resolved in this study.

T cell depletion outcomes and CR vs. non-CR-associated peripheral T cell signatures support a central requirement for CD8^+^ and CD4^+^ T cells in durable protection, consistent with a model in which a sufficient magnitude of effector T cell activity is germane for long-term protection. Furthermore, our longitudinal immunoprofiling findings rationalize CD40-licensed cDC subsets in the TDLN as potentially critical supporters of the observed T-cell-dependent tumor control. Future studies will directly test cDC subset requirements (e.g. cDC1 dependence) and interrogate implicated pathways such as type I interferon signaling and the CD70– CD27 axis^60,61^. Another plausible alternative hypothesis is that combinatorial efficacy resulted from reprogramming of MDSCs into pro-inflammatory monoctyes^62–64^. While we observed no changes in the number or percentage of intratumoral Arginine-1^+^ monocytic myeloid-derived suppressor cells (M-MDSCs) or G-MDSCs (Supplementary Fig. 6A-D), their frequency at a single time point may not fully capture myeloid functional state.

Our study aligns with others that have cemented the capacity of T-FUS to elicit immune-cooperative efficacy in BC^11,18,19^. However, no studies to date have combined a thermally ablative mode of FUS with CD40 agonism, nor have studies to date investigated FUS in any form with CD40 agonism in BC. Sub-ablative FUS heating has been paired with CD40 agonism as an ‘in situ vaccination’ strategy in melanoma. Indeed, in a bilateral B16F10 model, multiple sessions of mild FUS hyperthermia combined with repeat intratumoral αCD40 conferred moderate abscopal tumor control, elevated intratumoral T cell responses, and macrophage repolarization^65^. A subsequent study using mechanical FUS ablation (boiling histotripsy; BH) in murine melanomas similarly supported that a single session of BH-induced tumor disruption can cooperate with intratumoral αCD40 to amplify inflammatory and effector programs yielding tumor control as well as sensitize tumors to dual immune checkpoint blockade^66^ (ICB). Our study complements and expands this literature by showing that thermal ablation cooperates with CD40 agonism in the context of (i) a distinct solid tumor type and (ii) a less invasive αCD40 administration route (systemic). Our study is also distinct in its implementation of an immunotherapy priming regimen as opposed to concomitant treatment. In this regard, our findings are consistent with other studies supporting the benefit of neoadjuvant staging of immunotherapy prior to T-FUS^18,19^ or other interventions^67,68^.

The degree to which FUS-mediated injury productively interfaces with CD40 agonism appears context- and regimen-dependent. Indeed, recent work in a pancreatic cancer model demonstrated that a single BH treatment liberated tumor antigen locally but did not promote its appearance in the draining node or activate conventional DCs, nor did it enhance the activity of systemic αCD40^69^ - underscoring that factors such as ablation modality, drug dose, ablation coverage, and therapeutic sequencing may also dictate the coupling between innate immune priming and robust adaptive response driven by FUS and CD40 agonism. Where we observed intragroup heterogeneity within the same model and treatment arm, chronic immune profiling unveiled that TDLN T cell landscapes diverged by response class. We postulate that this may reflect potential differences in residual antigen burden subsequent to T-FUS as well as the state of immune resolution versus ongoing disease. This heterogeneity is clinically relevant and motivates identification of actionable variables and early biomarkers that stratify non-responders such that they can be converted or rescued through rationally layered strategies such as ICB^48,66^.

Given the surge of clinical trials evaluating FUS in combination with immunotherapies (including those ongoing in BC: NCT04796220, NCT03237572), these findings bear near-term translational relevance for developing an incisionless, immunologically active strategy with robustness across BC subtypes. Multiple clinical FUS platforms provide feasible paths to implementation^12^, and the clinical landscape for CD40 agonism is rapidly evolving, including the surge in Fc-engineered agents designed to improve therapeutic index^31,70,71^. A timely next step will be to test whether subtotal T-FUS similarly potentiates next-generation CD40 agonists that offer improved safety profiles. Our findings position the clinically ripe combination of subtotal T-FUS and CD40 agonism as a non-invasive strategy for eliciting durable immunological protection and provide an early mechanistic framework to extend its relevance within and beyond BC.

## METHODS

### Cell line and animal maintenance

E0771 and EMT6 cells were cultured in Dulbecco’s Modified Eagle’s Medium (DMEM, Gibco #11965-092) with 10% Fetal Bovine Serum (FBS, Gibco #11875-093). 4T1 cells were cultured in RPMI 1640 (+ L glut, Gibco #11875-093) supplemented with 10% FBS. BPRKP110 mouse mammary cancer cell line was obtained from Dr. Melanie Rutkowski, University of Virginia. BRPKP110 cells were maintained in RMPI with 10% FBS, 1X L-glutamine (Gibco #25030-081), 1mM sodium pyruvate (100 mM, Gibco #11360-070), 50μM beta-mercaptoethanol (Gibco, #21985023). Cells were maintained at 37ºC and 5% *CO*_2_ . Cells were thawed up to a maximum of three passages and maintained in logarithmic growth phase for inoculations. Prior to expansion and freezing, all cell lines were authenticated and confirmed to be mycoplasma-free.

All animal work was performed under a protocol approved by the Animal Care and Use Committee at the University of Virginia and conformed to the National Institutes of Health guidelines for the use of animals in research. Eight-to ten-week-old female C57Bl/6 or BALB/c mice were obtained from The Jackson Laboratory (Jax #000664 & # 000651, respectively). Mice were inoculated subcutaneously (s.c.) inoculated with E0771 or 4T1 cells (4 × 10□) or in the 4th mammary fat pad with BRPKP110 (5 × 10□) or EMT6 cells (4 × 10□). The administration volume was 100□µL per mouse. Mice were housed on a 12-h/12-h light/dark cycle and food was available to them at all times. Tumor growth was monitored daily with precision digital caliper measurements, and with B-mode ultrasound imaging at selected timepoints using the VEVO 2100 MS400. Digital caliper measurements were used to calculate tumor volume as follows: *volume* = (*length* × *width*^2^ )/2. When tumors reach approximately 60*mm*^3^, 28 days (E0771), 16 days (BRPKP110), 10 days (EMT6), and 11 days (4T1) post-inoculation, mice were randomized with in-house MATLAB code into experimental groups on the basis of pre-treatment tumor volume.

### Contralateral tumor rechallenge

E0771 tumor-bearing mice that experienced complete rejection of primary tumors following T-FUS+αCD40 treatment were rechallenged on the contralateral flank. Complete responders were inoculated with a 2.5x dose of (1 × 10^6^ E0771 tumor cells) approximately 150 days after tumor eradication. Age-matched naïve control mice were simultaneously challenged alongside the complete responders, serving as controls. Mice were housed and tumor growth was monitored as described above.

### Ultrasound tumor measurements and analysis

While under inhaled isoflurane anesthesia, mice were placed on the Vevo Imaging Station (VisualSonics) equipped with motor for automatic image acquisition. Tumors were covered with degassed ultrasound gel prior to 3D ultrasound imaging, acquired using 30 MHz MicroScan linear ultrasound imaging array (MS400; Sonics). Tumors were segmented in 3D Slicer^72,73^ by manually painting approximately every 10th slice followed by automatic filling between slices. Segment statistics were then calculated for each segmented tumor and volumes were obtained.

### Anti-CD40 therapy

For anti-CD40 therapy, rat anti-mouse agonistic CD40 antibody (*α*CD40; 500*μ*g/kg in 100*μ*L; clone FGK4.5; Ichorbio) was diluted in sterilized 1X PBS and administered i.p. every 3 days for a total of five doses. aCD40 treatment was initiated once tumors reached approximately 60*mm*^3^. Mice that did not receive *α*CD40 received an i.p. injection of equivalently dosed rat IgG2a isotype control antibody (Ichorbio) at analogous time points. For all therapeutic efficacy studies, control group received IgG2a isotype control and underwent ‘sham’ treatment as described below.

### *In vivo* ultrasound-guided FUS partial thermal ablation

Mice were treated with T-FUS seven days after the initial dose of αCD40, on day 35 (E0771), day 23 (BRPKP110), day 17 (EMT6), or day 18 (4T1 and 4T1-ZSGreen) post-implantation. On treatment day, mice were anesthetized with an i.p. injection of ketamine (50mg/kg; Zoetis) and dexmedetomidine hydrochloride (0.25mg/kg; Dechra). Tumors and surrounding region were depilated prior to T-FUS treatment, following which mice were positioned and treated. T-FUS was performed with a custom-built ultrasound-guided FUS system comprised of four 3.78 MHz single-element transducers (SU-102, Sonic Concepts), each with a diameter of 33 mm and a radius of curvature of 55 mm. The system was powered by a 200W amplifier (Electronics & Innovation 1020L) driven by an arbitrary function generator (Tektronix AFG3022C), and axially co-registered to a 15 MHz MicroScan linear ultrasound imaging array (MS200; VisualSonics). Degassed deionized water, maintained at 37ºC, was used as the acoustic coupling medium. A motorized 3D motion stage was used to adjust the position of the mouse relative to fixed transducer positions for each sonication. B-mode ultrasound imaging (VEVO 2100 MS200, VisualSonics) was employed for tumor identification and sonication planning. Sonication points were arranged within the ultrasound-visible tumor margins across a single plane of the tumor, with each point separated by 1.5 mm. T-FUS was applied continuously for 15 s at 18 W acoustic power over 2-4 planes of treatment, each of which was separated by 1 mm. We deployed a grading system that assigned blinded scores of 1-3 to the visible degree of hyperechoic signal across treatment planes in the four preclinical models of BC that underwent T-FUS (E0771, BRPKP110, EMT6, 4T1). Scoring was based on the degree of hyperechoic signal across each treated plane. A score of 1 indicated minimal hyperechogenicity confined to a single plane and not present throughout the entire treated area of that plane. A score of 2 represented a moderate level of hyperechogenicity across 2-3 planes, while a score of 3 reflected a high degree of hyperechogenicity observed across all planes (3-4) and in the majority or entirety of the treated areas within those planes. Mice that did not receive T-FUS treatment underwent “sham” treatment, consisting of anesthesia, depilation, and analogous partial exposure to the 37ºC degassed water bath for 3 minutes. At the conclusion of “sham” or T-FUS treatment, mice were transferred to a heating pad, given Antisedan for anesthesia reversal, and recovered.

### T cell depletion

E0771-bearing C57Bl/6 mice underwent depletion of CD4□ T cells, CD8□ T cells, or both 1 day following conclusion of T-FUS+*α*CD40 dosing paradigm (day 41). T cell depletion was carried out with anti-CD8 (*α*CD8; 2.43 clone; Bio X Cell) and anti-CD4 (*α*CD4; GK1.5 clone; Bio X Cell). Depleting antibodies were administered intravenously (i.v.) every 3 days for a total of seven doses. To confirm depletions, tail vein bleeds were performed on day 46 and circulating CD4□ and CD8□ T cell levels were screened via flow cytometry.

### Immunohistochemistry

On day 36 (E0771), IgG-control or T-FUS treated tumors underwent cardiac perfusion with a sequence of 1X PBS, 10% neutral buffered formalin (NBF), 1X PBS. 24 hours later, fixed tumors were paraffin embedded, sectioned, and stained for hematoxylin and eosin (H&E), cleaved caspase 3 (CL Casp. 3; Cell Signaling, Cat #9661), poly-ADP ribose polymerase 1 (PARP1; Abcam, ab32064), and heat shock protein 70 (HSP70; Abcam, Clone: EPR16892, ab181606). Digital scans of stained slides were collected using ZEISS Axioscan 7. Demarcation of ablative and periablative zone was determined by evaluating margins of apoptosis signatures identified by H&E, CL Casp3, and PARP staining as well as immunomodulatory changes indicated by HSP70 expression.

### *In vitro* and *in vivo* ATP analysis

Luciferin-luciferase optical imaging was performed to quantify ATP production following T-FUS treatment. *In vitro*, 4T1 cells were washed with 1X PBS prior to being prepared in 60uL serum-free, phenol-red-free RPMI and suspended in PCR tubes at 3x10^6^ cells per tube. Immediately following ablation of 4T1 cells *in vitro*, supernatants were collected for ATP quantification using the CellTiter-Glo® Luminescent Cell Viability Assay (Promega G7571). Luminescence was measured with a TECAN Spark Multimode Microplate Reader.

*In vivo* ATP production was evaluated in E0771 tumors following T-FUS treatment. Mice were injected intravenously with 3mg D-luciferin (Invitrogen, L2916) 1 minute prior to T-FUS treatment. Firefly luciferase (Biotium #30020-1) was administered immediately after the conclusion of T-FUS treatment. Bioluminescence imaging (Lago X) was performed over 10-40 minutes post-treatment. Tumor-localized luminescence was quantified using Aura Imaging Software (Spectral Instruments Imaging).

### Flow cytometry

Tumor-bearing mice were euthanized at various time points for tissue and/or blood collection and subsequent immunoprofiling by multispectral flow cytometry. In E0771 tumor harvests, to distinguish between tissue-resident and vascular immune cell populations, mice were injected i.v. with rat anti-mouse CD45 PerCP (clone 30-F11; BioLegend) 3 mins before euthanasia. Tumors, spleens, inguinal tumor draining lymph nodes (TDLNs), and contralateral nondraining inguinal lymph nodes (cLNs) were mechanically homogenized with glass homogenizers and filtered through 100*μ*m nylon filter mesh to obtain single-cell suspensions. Tumors went through an additional enzymatic digestion step prior to filtration with collagenase (Gibco #17018-029) and DNase 1 (Roche #10104159001). Following filtration, lymphocyte layer was collected subsequent to isolation procedure using Lympholyte-M Cell Separation Media (Cedarlane #CL5035). Spleen and blood samples were also subjected to hemolysis using BioLegend 1X Red Blood Cell (RBC) lysis buffer (BioLegend #420301). Following RBC lysis, samples were quenched with complete media (RPMI or DMEM + 10% FBS).

Following, cells were centrifuged at 1500 RPM for 5 minutes, washed with 1X PBS, and centrifuged again at the same parameters. Cells were stained with either Fixable Live/Dead Aqua or Near IR for 30 mins at 4ºC. Following, cells were fixed with anti-mouse CD16/32 (ThermoFisher #14-0161-86) to block Fc gamma receptors for 15 mins at 4ºC, then centrifuged and washed twice with 1X Flow Cytometry Staining Buffer (FACS; R&D Systems #FC001). Surface staining was performed in Brilliant Stain Buffer (BD Biosciences #566349) for 30 mins at 4ºC with fluorescent monoclonal antibodies for CD11b, MHCII, CD317, F4/80, Ly6C, Ly6G, XCR1, CD19, CD40, CD11c, CD86, SIRP*α*, CD45, CD45.2, CD3, DX5, CD8, CD4, TIM3, LAG3, PD1, PDL1, CD103, CD69, CD44, CD62L, CD40L, and CD25. Intracellular staining was carried out with Foxp3/Transcription Factor Staining Buffer Set (Tonbo Biosciences #TNB-0607-KIT). Samples were additionally stained for iNOS, Ki-67, Arg1, CD206, TNF-α, Granzyme B, TOX, IFN-γ, FOXP3.

Table Supplementary 2 presents comprehensive details on all fluorescently labeled antibodies, including their clones and vendor. Samples were fixed with 1X FACS Lysis Solution (BD Biosciences #34902) for 10 mins at 4ºC, then centrifuged and resuspended with FACS buffer. Flow cytometry was carried out using the Cytek Aurora Borealis 5 laser and SpectroFlo v3.0.3 software. Gating and data analysis was performed using FlowJo 10 or OMIQ software. Representative gating strategies are depicted in Supplementary Figures 7-8.

### Ex Vivo T Cell Stimulation

On day 7 post-E0771 tumor rechallenge, blood samples were prepared as described above. Cells were plated in two 96-well round-bottom plates: one stimulated and the other non-stimulated. Stimulated plates received the Cell Stimulation Cocktail (eBioscience™ #00497503) comprised of phorbol 12-myristate 13-acetate (PMA), ionomycin, brefeldin A and monensin, while non-stimulated cells were plated in media. Plates were maintained in 37ºC and 5% *CO*_2_ for 2.5 hours. Cells were then stained for surface and intracellular markers (Ki-67, granzyme B, IFN-γ, and TNF-α).

### Brefeldin A Injection

To inhibit the secretion of cytokines from cells, E0771 tumor-bearing mice were treated with Brefeldin A (BFA; 250 μg; Selleck Chemicals #S7046) diluted in sterilized 0.9% saline. BFA was administered i.p. to mice 4 hours prior to harvest.

### Dimensional Reduction Analysis of Flow Cytometry Data

OMIQ software (Dotmatics; www.omiq.qi) was used to perform dimensionality reduction using t-distributed stochastic neighbor embedding (tSNE) of the flow cytometry data. Briefly, gating strategies from FlowJo were imported into OMIQ to maintain consistent gating on CD45.2□ or CD45□ live cells by removing cell debris, doublets, and dead cells. All fluorescent parameters – excluding viability dye Blue or Aqua, CD45, and/or CD45.2 – were included as dimensions. Subsampling was performed with 20,000 events. Single live CD45 cells for each file were concatenated for analysis by tSNE dimensionality reduction followed by multicolor mapping of each group in the concatenated file to identify proportions of each population.

### Statistical Analysis

Statistical analyses were performed using GraphPad Prism 10.1.0. A comprehensive overview of the statistical techniques used in each experiment can be found in the accompanying figure legend. “n.s.” denotes that data are not significant. All data are represented as mean ± SEM. Normality was assumed where no significant deviation was detected by Q-Q plots and the Shapiro-Wilk test. Equal variance was not assumed where homoscedasticity plot and Bartlett’s test indicated significant deviation. Therefore, Welch’s ANOVA or Welch’s t-test was employed. For multiple comparisons, pairwise analyses were performed against the T-FUS+αCD40 group, and Dunnett’s test was used to adjust for multiple comparisons.

## Supporting information

Demir-FUSCD40-Supplement

## Acknowledgments

We thank the Flow Cytometry Core Facility of the University of Virginia for their support and use of their cytometers, supported by the National Cancer Institute P30-CA044579 Center Grant. This work was supported by Biorepository and Tissue Research Facility which is supported by the University of Virginia School of Medicine, Research Resource Identifiers (RRID): SCR_022971. This work used FFPE embedded and sectioning in the Research Histology Core Facility which is supported by the University of Virginia School of Medicine, Research Resource Identifiers (RRID): SCR_025470. We thank Dr. Drew Dudley (UVA) for providing the E0771 cell line. We thank Dr. Frederic Padilla for his expertise and development of the ultrasound-guided FUS system used herein. We thank Drs. Patrick Dillon and David Brenin (UVA) for their invaluable clinical insights throughout the development of this work. This research was supported by state funding within the University of Virginia Comprehensive Cancer Center.

## Author Contributions

Conceptualization: ZEFD, NDS

Methodology: ZEFD, AK, MRD, NDS

Formal Analysis: ZEFD, AK, ML

Investigation: ZEFD, BGA, ML, TS, SM, ATT, MRD

Writing - Original Draft: ZEFD

Writing - Review and Editing: ZEFD, AK, MRD, MR, NDS

Resources: MR

Funding acquisition: NDS

Supervision: NDS

