## Supplementary material for "Focused Ultrasound Thermal Ablation and CD40 Agonism Reprograms Breast Tumor Immunity to Drive Regression and Memory": Demir-FUSCD40-Supplement

**Table 1: Percentage of each grading scores for each model – E0771, BRPKP110, EMT6 and 4T1 – and collectively across all four breast cancer models.**

| Tumor-bearing Model | Grade 3 | Grade 2 | Grade 1 |
|---|---|---|---|
| E0771 | 78.6% | 14.3% | 7.1% |
| BRPKP110 | 88.9% | 11.1% | 0% |
| EMT6 | 31.3% | 56.3% | 12.5% |
| 4T1 | 94.4% | 0% | 5.6% |
| **Overall Percentage** | 73.2% | 18.3% | 8.5% |

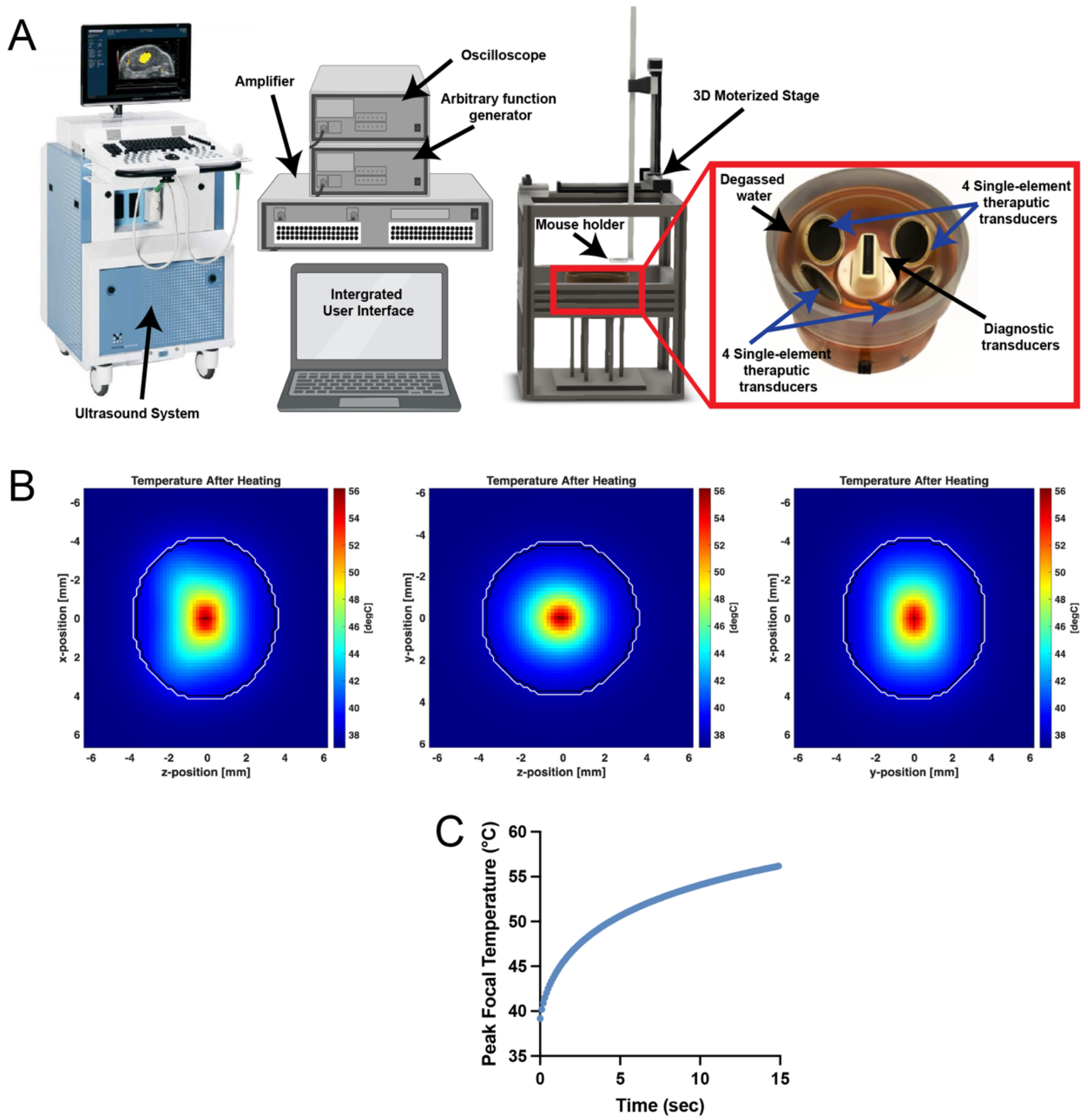


**Supplementary Figure 1: 3D computational model reveals clear rise in temperature and ablation volume following T-FUS treatment.** A) Illustration of custom ultrasound-guided focused ultrasound system. B) In silico model rendering temperature distributions after T-FUS treatment in xz, yz, and xy planes through the tumor center. C) Peak focal temperature achieved in the tumor throughout the T-FUS treatment.

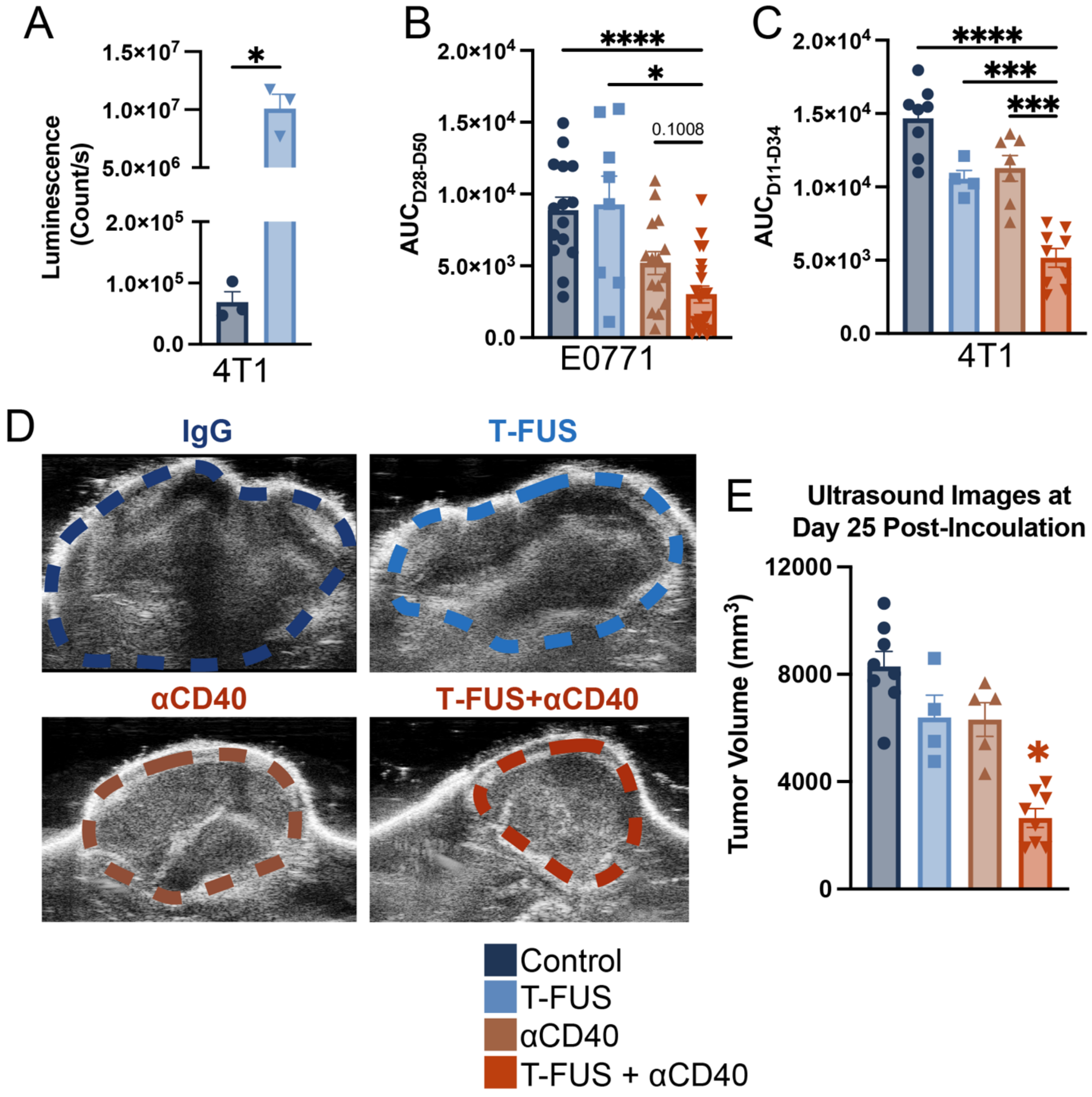


**Supplementary Figure 2: Thermal ablation enhances ATP release and, in combination with αCD40, constraints tumor outgrowth.** (A) Elevated average luminescence of ATP released from 4T1 cells in vitro immediately following T-FUS. Significance assessed by Welch's T-test.*p < 0.05, **p < 0.01 vs. control. B-C) Area under the curve (AUC) analysis of E0771 and 4T1 tumors. (D) Representative ultrasound images depicting degree of 4T1 tumor-burden. (E) Tumor volumes of 4T1 tumor-bearing mice at day 25 post-inoculation, assessed by ultrasound imaging. Significance assessed using a Welch ANOVA. *p < 0.05, **p < 0.01 vs. T-FUS+αCD40.

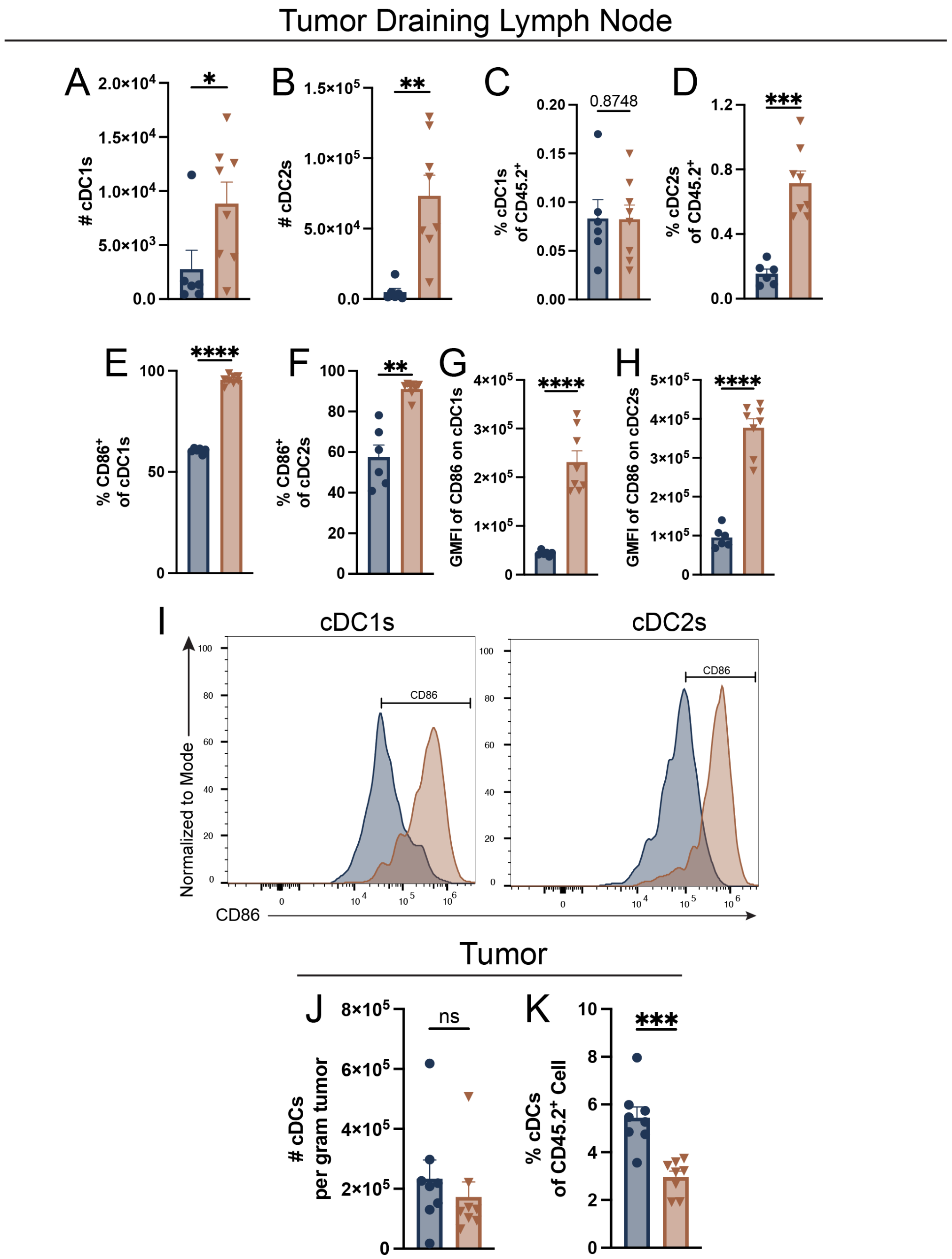


**Supplementary Figure 3: aCD40 priming prior to T-FUS activates cDC1 and cDC2 in the TDLN.** (A-B) Absolute number and (C-D) percentage of cDC1s and cDC2s in the TDLN. (E-F) Proportion of $CD86^+$ cDC1s and cDC2s in the TDLN. (G-H) The GMFI of CD86 on cDC1s and cDC2s in the TDLN. (I) Histogram representing the percentage of $CD86^+$ cDC1s and cDC2s in the TDLN. J) Absolute number of cDCs per gram of tumor and K) percentage of cDCs in the tumor. Significance assessed by Welch's T-test.*$p < 0.05$, **$p < 0.01$ vs. control.

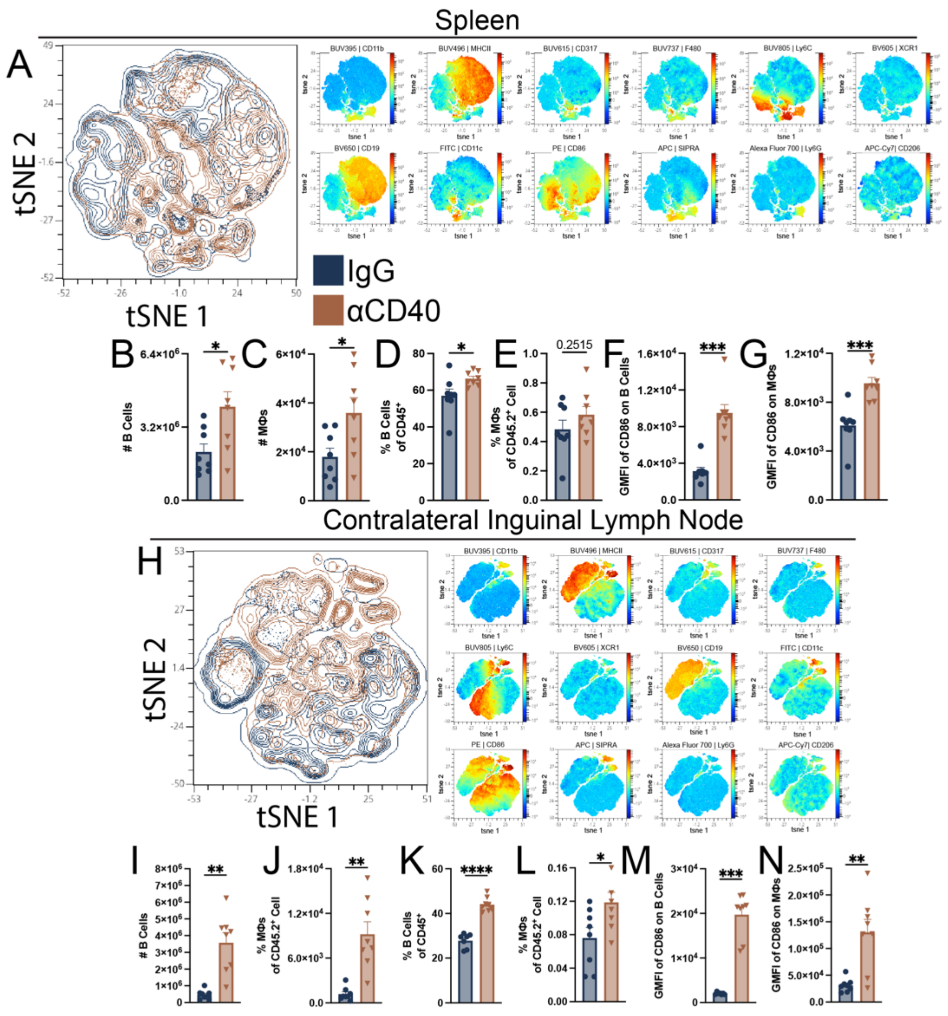


**Supplementary Figure 4: Priming with a CD40 agonist promotes systemic activation of APCs.** (A) Multigraph color mapping of tSNE plot on $CD45^+$ cells in the spleen. (B-C) Absolute number and (D-E) percentage of circulating B cells and MΦs in the spleen. (F-G) The GMFI of CD86 on B cells and MΦs in the spleen. (H) Multigraph color mapping of tSNE plot on $CD45^+$ cells in the contralateral lymph node. (I-J) Absolute number and (K-L) percentage of circulating B cells and MΦs in the contralateral lymph node. (M-N) The GMFI of CD86 on B cells and MΦs in the contralateral lymph node. Significance assessed by Welch's T-test.*$p < 0.05$, **$p < 0.01$ vs. control.

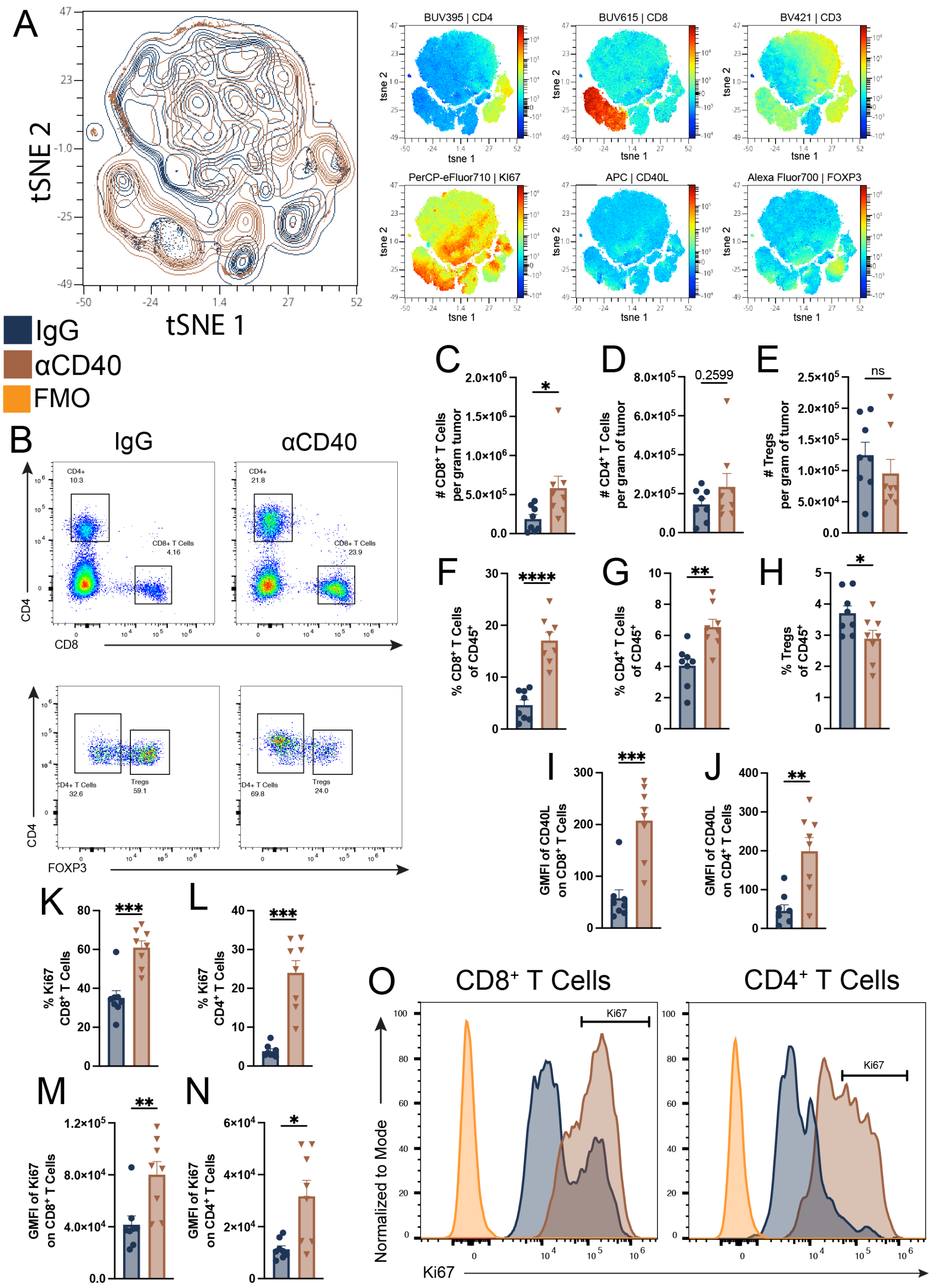


**Supplementary Figure 5: CD40 priming promotes intratumoral T lymphocyte population.** (A) Multigraph color mapping of tSNE plot on CD45$^+$ cells in the tumor. (B) Representative dot plot shows changes in CD8$^+$ and CD4$^+$ T cells population in the tumor. (C-E) Absolute number and (F-H) percentage of intratumoral CD8$^+$ T cells, CD4$^+$ T cells, and Tregs. (I-J) The GMFI of CD40L on CD8$^+$ and CD4$^+$ T cells population. (K-L) Proportion of Ki67$^+$ CD8$^+$ and CD4$^+$ T cells in the tumor. (M-N) The GMFI of Ki67 on CD8$^+$ and CD4$^+$ T cells. (O) Representative dot plot of intratumoral CD8$^+$ and CD4$^+$ T cells. Significance assessed by Welch's T-test.*$p < 0.05$, **$p < 0.01$ vs. control.

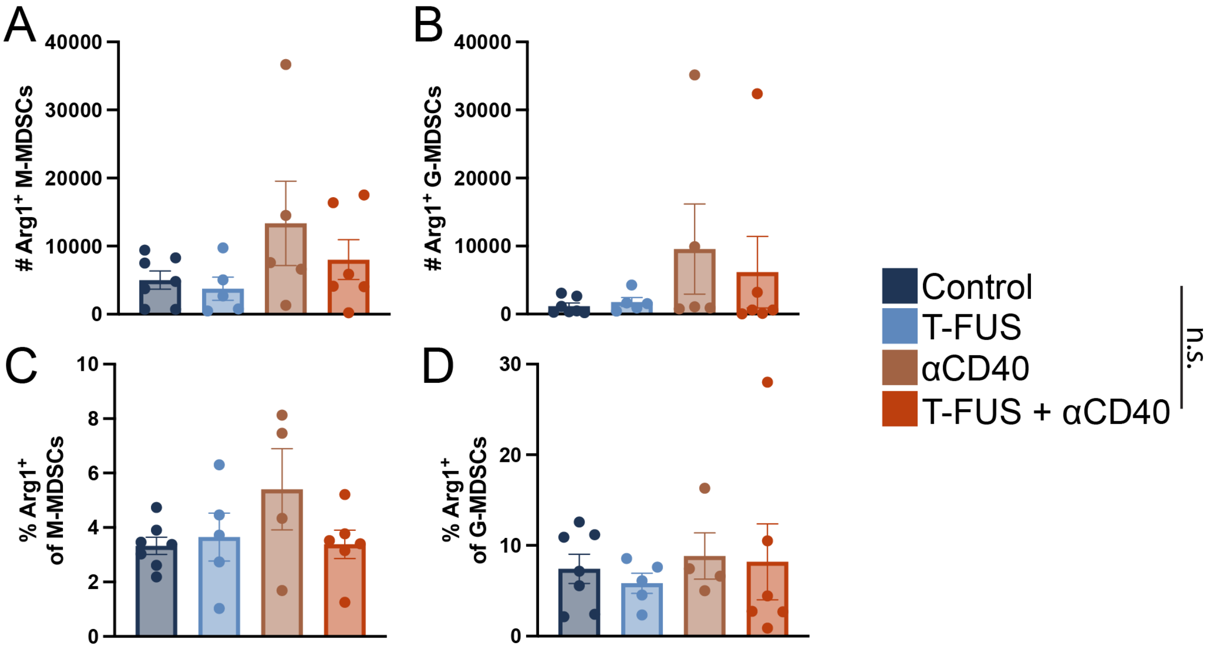


**Supplementary Figure 6: T-FUS with αCD40 does not alter intratumoral Arg1-positive M-MDSCs or G-MDSCs population.** (A-B) Absolute number and (C-D) percentage of intratumoral Arg1$^{+}$ M-MDSCs or G-MDSCs. Significance assessed using a Welch ANOVA. *$p < 0.05$, **$p < 0.01$ vs. T-FUS+αCD40.

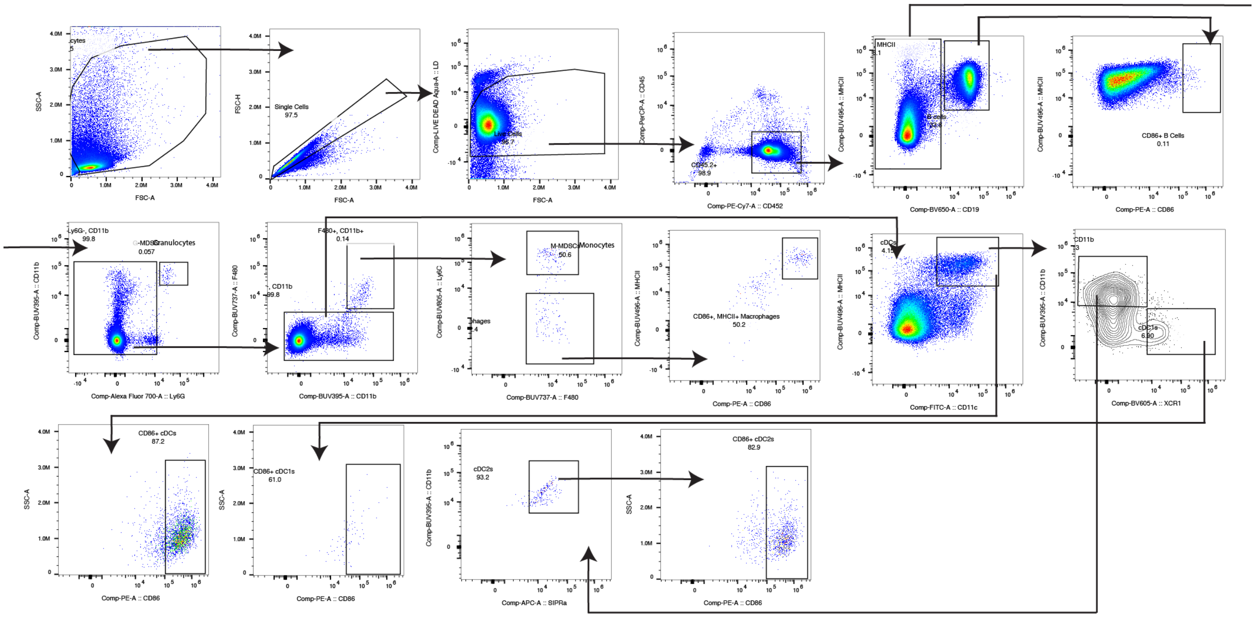

**Supplementary Figure 7: Representative flow cytometry gating strategy used to identify antigen presenting cells (APCs).**

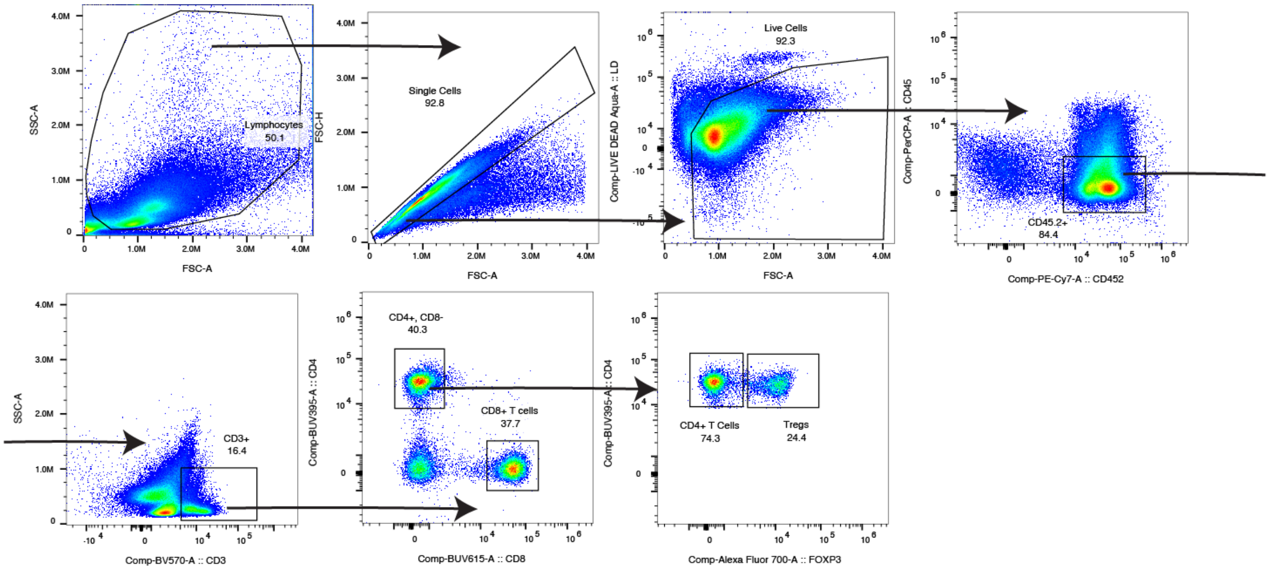


**Supplementary Figure 8: Representative flow cytometry gating strategy used to identify T lymphocytes.**
